# tRNA-Based Polycistronic CRISPR/Cas9 System Boosts Efficiency of Multi-Gene Deletion in the Moss Physcomitrella

**DOI:** 10.1101/2025.07.29.667574

**Authors:** Elena Kozgunova

## Abstract

CRISPR/Cas9-based genome editing in the model bryophyte *Physcomitrium patens* (commonly known as Physcomitrella) is widely used for gene knockout via small insertions or deletions (indels). However, this approach may leave residual gene activity and typically requires sequencing-based validation. In this study, we established an efficient strategy for generating large, targeted deletions across multiple genes using dual-gRNA targeting. We first compared the efficiency of polycistronic tRNA-gRNA arrays to conventional gRNA constructs expressed under individual promoters, using the checkpoint protein gene *MAD2* as a target. We found that a polycistronic construct doubled the frequency of large gene deletions compared to a conventional design. We then demonstrated that simultaneous deletion of two or four genes, targeting the *katanin* and *TPX2* gene families, respectively, can be achieved in a single transformation event. The polycistronic system also increased deletion frequencies in the multiplex context, with up to 42% efficiency for individual genes and successful recovery of quadruple mutants. As a drawback, we confirmed that deletion efficiency varied substantially among individual gRNA pairs, indicating that gRNA design remains a critical factor in multiplex editing. This study establishes a versatile and scalable framework for generating multi-gene deletion mutants in *P. patens*, facilitating functional genomics and biotechnological applications requiring precise gene removal.

## Introduction

The CRISPR-Cas9 system has revolutionized genome editing by enabling precise genetic modifications. It is widely used to introduce frameshift mutations by generating small insertions or deletions (indels) at target sites, leading to gene knockout through the disruption of the reading frame (Jiang and Doudna, 2017; Adli, 2018). However, this approach has limitations: mutation verification requires sequencing, and in some cases, truncated proteins may still be expressed from downstream start codons, potentially retaining partial gene function (Smits et al., 2019; Hosur et al., 2020).

To overcome these limitations, targeting a gene with two guide RNAs (gRNAs) can induce large deletions, removing most or all of the gene coding sequence (Canver et al., 2014; Chen et al., 2014; Zhou et al., 2014). This approach offers advantages, such as straightforward detection by genotyping PCR and gel electrophoresis, as well as eliminating the risk of truncated protein expression. On the other hand, the requirement for multiple gRNAs per gene can potentially increase the risk of off-target effects (Chen et al., 2014; Zhou et al., 2014).

The moss *Physcomitrium patens,* commonly known as Physcomitrella, is recognized as a powerful model organism for evolutionary and developmental (evo-devo) studies due to its simple genome and ease of genetic manipulation (Rensing et al., 2020). Beyond fundamental research, *P. patens* has emerged as a robust platform for biopharmaceutical production, where genome editing is required to engineer metabolic pathways and optimize target yields (Decker and Reski, 2020; Horn et al., 2021).

The introduction of CRISPR/Cas9 in *P. patens* provided a versatile, efficient, and scalable tool for genome editing (Lopez-Obando et al., 2016; Nomura et al., 2016). Since then, CRISPR applications in *P. patens* have rapidly evolved, with modular vector systems enhancing editing flexibility (Mallett et al., 2019) and multiplex mutagenesis enabling simultaneous targeting of complex gene families, such as phytochromes (Trogu et al., 2021). CRISPR/Cas9 in *P. patens* can be used not only for introducing mutations through double-stranded breaks, but also for fine-tuned gene editing. For instance, transient co-transformation strategies combining CRISPR/Cas9 with oligonucleotide templates were demonstrated to facilitate precise template insertions (Yi and Goshima, 2020). Techniques such as prime editing (Perroud et al., 2022, 2023) and base editing (Guyon□Debast et al., 2021) were developed for gene editing without introducing double-strand breaks. Alternatives to CRISPR/Cas9, such as CRISPR/LbCas12a, were also developed for P. patens genome editing with demonstrated high efficiency in multiplex editing (Pu et al., 2019). Studies also highlight the role of DNA repair pathways in CRISPR outcomes, with POLQ identified as a key player in double-strand break repair (Mara et al., 2019). In addition to targeted knockouts, CRISPR/Cas9 in *P. patens* was also recently utilized for genetic screening to identify novel cell division genes (Maekawa et al., 2025).

In this study, we aimed to simultaneously introduce large deletions into multiple genes in *P. patens* via CRISPR/Cas9, removing most of the coding sequences. Notably, large deletions within single genes, ranging from 3000 to 6000 bp, have already been demonstrated as feasible using CRISPR/Cas9 in *P. patens* (Nomura et al., 2016; Collonnier et al., 2017; Trogu et al., 2021). Building upon these findings, we specifically investigated whether similar large-scale deletions could be extended to multiple genes in a single transformation event.

In this study, we tested several CRISPR/Cas9-based strategies to induce large deletions across multiple gene loci in *P. patens*. We systematically compared conventional and polycistronic gRNA constructs, evaluated deletion efficiencies across different target genes, and examined whether simultaneous removal of coding regions from multi-gene families could be achieved in a single transformation event. Using dual-gRNA targeting, we achieved large deletions at four independent loci, ranging from ∼2.2 kb to ∼7.8 kb.

We also demonstrated that polycistronic constructs improved deletion efficiency by approximately 20% in both single- and multi-gene editing contexts. These results establish a scalable and efficient approach for multiplex gene knockout in *P. patens*, with applications in functional genomics and genome engineering.

## Results

### Polycistronic gRNA expression increases gene deletion efficiency

In this study, we tested two main hypotheses: (1) whether increasing the number of guide RNAs (gRNAs) enhances the efficiency of large gene deletions, and (2) whether polycistronic gRNA constructs utilizing the transfer RNA(tRNA) processing strategy are functional in Physcomitrella.

To address the first hypothesis, we compared the gene deletion efficiency of two gRNAs targeting the start and end of the gene, versus deletion efficiency six-gRNA construct, three targeting the beginning and three targeting the end of the coding region. For the second hypothesis, we examined whether the tRNA strategy, which enables multiple gRNAs to be expressed under a single promoter by incorporating tRNA spacers, is effective in Physcomitrella. This strategy relies on endogenous RNA processing systems to cleave the tRNA-derived hairpins, thereby releasing individual gRNAs from a polycistronic transcript (Xie et al., 2015; Zhang et al., 2016).

These hypotheses were tested using the checkpoint protein gene *MAD2* (Pp3c4_13910V3.1) as a deletion target. *MAD2* was reported to be a non-essential gene under normal growth conditions in both *P. patens* (Kozgunova et al., 2019) and *A. thaliana* (Komaki and Schnittger, 2017). Four types of plasmid constructs were designed:

- **Construct A**: Two gRNAs, each driven by its own U6 promoter, targeting the start and end of the *MAD2* gene.
- **Construct B**: Six gRNAs, each under the individual U6 promoter, with three targeting the start and three targeting the end of the *MAD2* gene.
- **Construct C**: A polycistronic construct driven by a single U6 promoter, containing two gRNAs separated by tRNA-derived hairpins (same gRNAs as Construct A).
- **Construct D**: A polycistronic construct under a single U6 promoter, containing six gRNAs separated by tRNA-derived hairpins (same gRNAs as Construct B) (Figure 1A).

**Figure 1.**
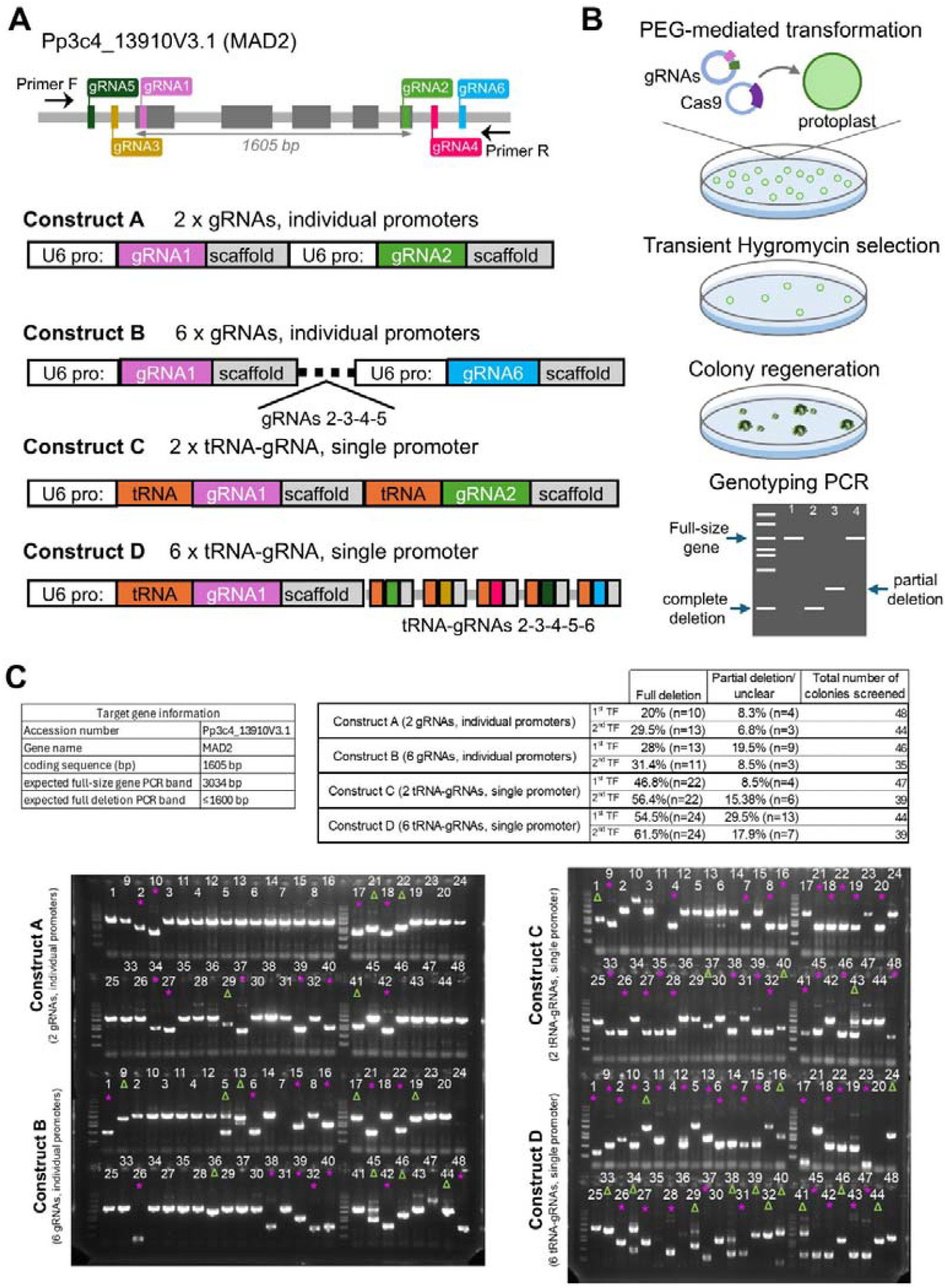
Polycistronic tRNA-based gRNA expression increases gene editing efficiency in P. patens. **(A)** Schematic representation of the MAD2 gene with gRNA target positions and four different gRNA expression cassettes. **(B)** Schematic illustration of the experimental workflow. **(C)** (Top) Summary of MAD2 deletion efficiencies using Constructs A, B, C, and D. (Bottom) Genotyping PCR results. Full deletion bands are marked with magenta asterisks; partial deletion or unclear results are indicated with green arrowheads.

For all experiments, Cas9 nuclease was supplied in a separate plasmid and co-transformed with the gRNA constructs into Physcomitrella protoplasts using the standard PEG-mediated transformation method (Yamada et al., 2016). The gRNA plasmids also carried a hygromycin resistance cassette, and transient selection pressure was applied for 5 to 7 days. Following colony regeneration, genotyping PCR was performed using primers flanking the target deletion region (Figure 1B).

All tested constructs successfully introduced large gene deletions into the *MAD2* gene. Notably, the polycistronic gRNA-tRNA construct (Construct C) demonstrated twice the efficiency of the conventional construct with individual U6 promoters (Construct A), with full gene deletion detected in 51.6±4.8% of colonies compared to 24.75±4.75% in Construct A (Figure 1C). Given that the same gRNA sequences were used in both constructs, the increase in gene deletion cannot be attributed to the different efficiency of gRNAs.

Introducing multiple gRNAs slightly improved the efficiency of large gene deletions but also resulted in a higher occurrence of partial gene deletions (Figure 1C). Full deletions were slightly more common in both conventional constructs (24.75±4.75% in Construct A and 29.7±1.7% in Construct B) and polycistronic constructs (51.6±4.8% in Construct C and 58±3.5% in Construct D). The multiple gRNAs and polycistronic constructs have also slightly increased the number of colonies with only partial deletions—7.55±0.75% for Construct A vs 14±5.5% for Construct B, 11.94±3.44% for Construct C and 23.7±5.8% for Construct D.

Considering that increasing the number of gRNAs did not provide a substantial improvement in full gene deletion efficiency and also comes with the elevated risk of off-target effects, we concluded that using two gRNAs per gene target is a more practical and sufficient strategy for future experiments.

### Modular-based CRISPR/Cas9 system

To enhance editing efficiency and streamline transformation workflows, we developed a modular CRISPR/Cas9 system that integrates all gRNA expression cassettes and the Cas9 nuclease into a single binary vector. As part of this optimization, we replaced the previously used rice actin 1 promoter (Lopez-Obando et al., 2016) with the 677 bp *lhcsr1* promoter (*Pp3c9_3440V3.1*), which exhibits high activity in *P. patens* protoplasts (Hiss et al., 2017).

Previously developed modular CRISPR/Cas9 system utilized Gateway-based cloning to incorporate up to 4 gRNA cassettes into the Cas9-containing vector (Mallett et al., 2019). Here, we developed a modular system that utilizes Golden Gate assembly for the scalable construction of gRNA arrays. We developed three versions of the system to accommodate 4, 6, or 8 gRNAs, each expressed under its own *PpU6* promoter and followed by a gRNA scaffold.

gRNAs are first cloned into entry vectors via BbsI sites and then assembled into the final Cas9-containing destination vector using BsaI-mediated Golden Gate cloning, guided by unique four-base overhangs to ensure ordered integration. The destination vector also carries Cas9 driven by the *lhcsr1* promoter, and a hygromycin resistance cassette under the CaMV 35S promoter for selection in moss (Figure 2A).

**Figure 2.**
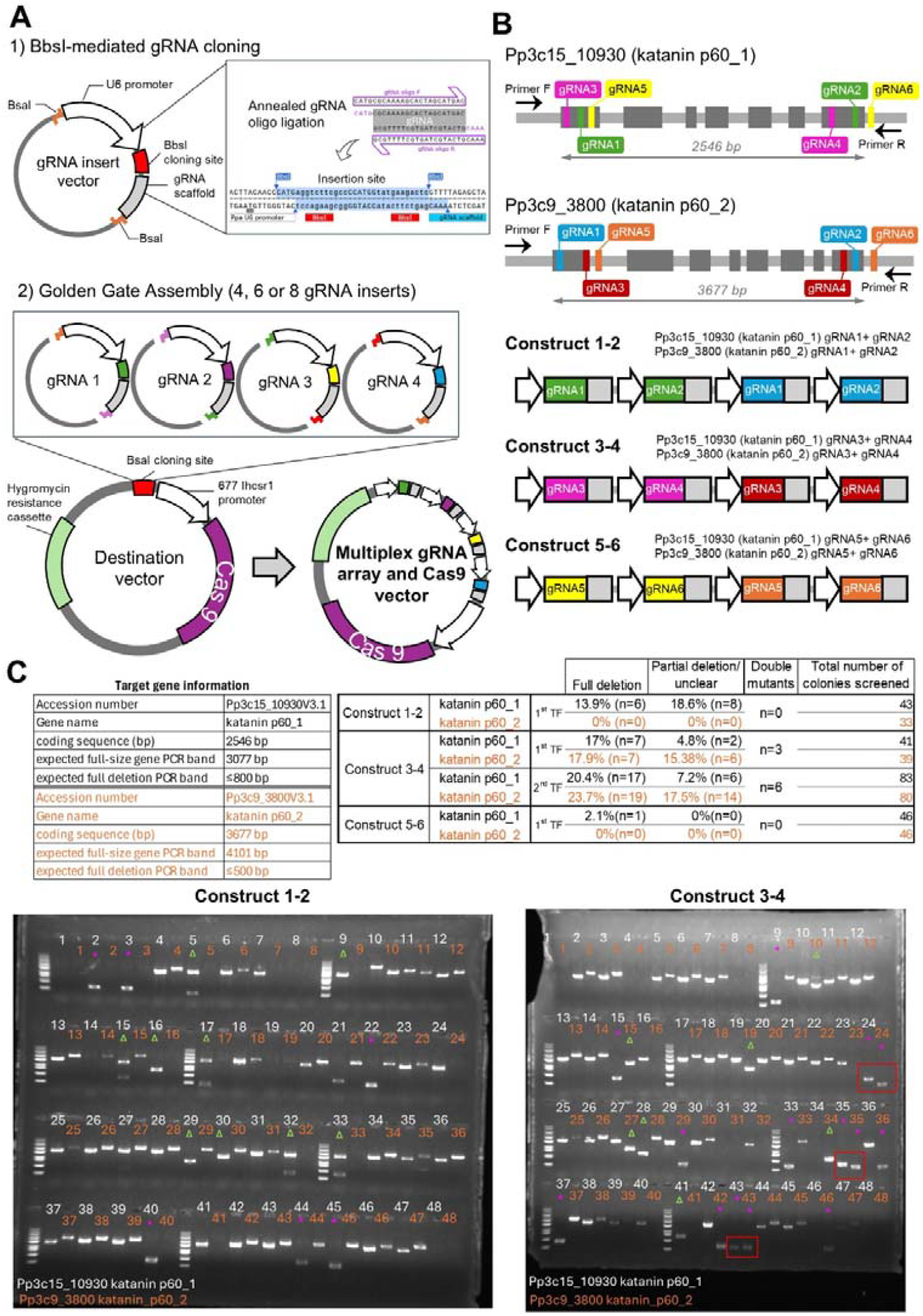
Golden Gate-based modular assembly for multiplex gene editing in P. patens. **(A)** Schematic illustration of the Golden Gate-based modular assembly **(B)** Schematic representation of the *katanin p60_1* and *katanin p60_2* genes with gRNA target positions and three gRNA expression cassettes. **(C)** (Top) Summary of *katanin p60_1* and *katanin p60_2* deletion efficiency using Constructs 1-2, 3-4, and 5-6. (Bottom) Genotyping PCR results of Construct 1-2 and 3-4. Full deletion bands are marked with magenta asterisks; partial deletion or unclear results are indicated with green arrowheads. Double mutants are indicated with red squares

### Simultaneous Deletion of Two *Katanin* Genes

Next, we attempted the simultaneous deletion of multiple genes. We selected the microtubule-associated protein *katanin*, which has two gene homologs and has been functionally characterized in *P. patens* (Nakaoka et al., 2015). For each gene, we designed two gRNAs, one targeting the 5′ region (typically the first exon) and one targeting the 3′ region (typically the last exon). The positions and sequences of these gRNAs are listed in Supplemental Table 1. In this experiment, each gRNA was expressed as an individual construct, driven by the U6 promoter and followed by the gRNA scaffold (Figure 2B).

In our initial attempt at simultaneous multi-gene deletion, results were limited. Large deletions were observed only in one gene, *katanin p60_1 (Pp3c15_10930V3.1)*, while *katanin p60_2 (Pp3c9_3800V3.1)* remained unaffected (Figure 2C, Construct 1-2). We hypothesized that this outcome may be attributed to suboptimal gRNA efficiency or limited chromatin accessibility at the targeted loci.

To test this hypothesis, we designed two new sets of gRNAs targeting the *katanin* genes (Figure 2B). The original multiplex gRNA vector was designated Construct 1-2, while the newly designed vectors were named Construct 3-4 and Construct 5-6. Each construct contained two distinct gRNAs pairs targeting the 5′ and 3′ regions of *katanin p60_1* and *katanin p60_2*, respectively (Figure 2B).

Following moss transformation and genotyping PCR, Construct 3-4 successfully generated multiple colonies with confirmed deletions in both *katanin* genes (Figure 2C). Full deletions of *katanin p60_1* were observed in 18.7□±□1.7% of screened plants, and *katanin p60_2* deletions were detected in 20.8□±□2.9%. In the first transformation, we isolated three moss plants with large deletions in both genes; in the second transformation, six double mutants were recovered. These findings highlight the critical importance of gRNA sequence design and target site accessibility in achieving efficient multiplex gene editing in *P. patens*.

### Generation of Quadruple Deletions of Four *TPX2* Genes in a Single Transformation Event

We next investigated whether it is feasible to generate quadruple full-gene deletion mutants in a single transformation experiment. To this end, we selected *TPX2* (*Targeting Protein for XKlp2*), a conserved microtubule-associated protein involved in spindle assembly and positioning. In *P. patens*, four homologs of *TPX2* have been functionally characterized and shown to play roles in spindle morphogenesis and asymmetric cell division (Kozgunova et al., 2022).

Given our prior observation that some gRNA pairs are inefficient at inducing large deletions, we designed three independent CRISPR constructs: Constructs 1-2, 3-4, and 5-6, each containing a unique set of gRNAs for each of the four *TPX2* homologs. The gene structures and gRNA target sites are illustrated in Figure 3A; all gRNA sequences are provided in Supplemental Table 1.

**Figure 3.**
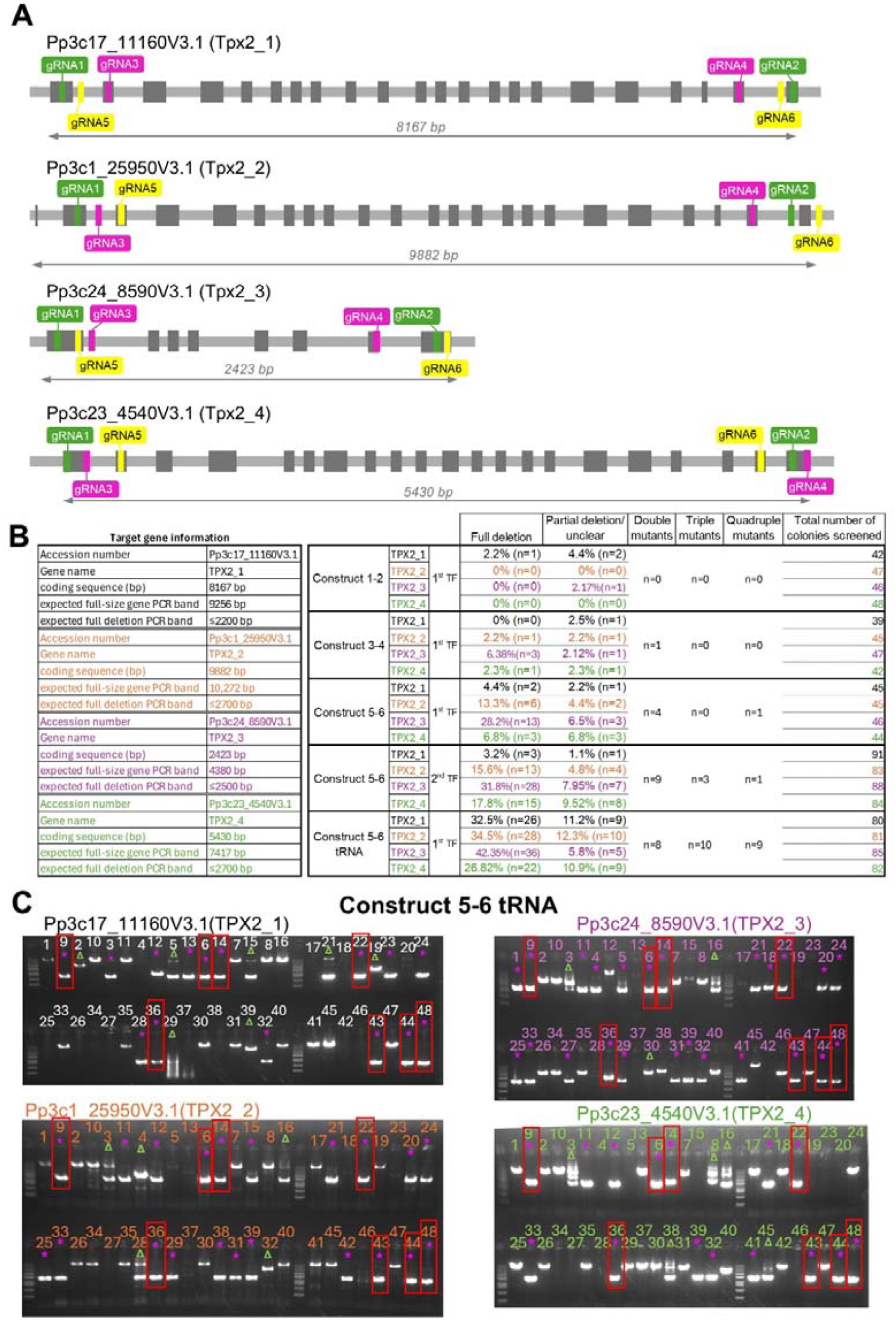
Quadruple TPX2 mutants with large deletions in four genes isolated in a single transformation. **(A)** Schematic representation of the *TPX2_1, TPX2_2, TPX2_3* and *TPX2_4* genes with gRNA target positions. **(B)** Summary of *TPX2s* deletion efficiency using Constructs 1-2, 3-4, 5-6, and polycistronic Construct tRNA 5-6. **(C)** Partial genotyping PCR results of Construct tRNA 5-6. Full deletion bands are marked with magenta asterisks; partial deletion or unclear results are indicated with green arrowheads. Quadruple mutants are indicated with red squares

Among the three designs, Construct 5–6 yielded the most promising results, with large deletions detected in all four *TPX2* genes, albeit with variable efficiency. Specifically, full deletions were observed in 3.8□±□0.6% of plants for *TPX2_1* (Pp3c17_11160V3.1), 14.45□±□1.15% for *TPX2_2* (Pp3c1_25950V3.1), 30□±□1.8% for *TPX2_3* (Pp3c24_8590V3.1), and 12.3□±□5.5% for *TPX2_4* (Pp3c23_4540V3.1).

Despite the relatively low efficiency for *TPX2_1*, we successfully isolated one quadruple mutant per transformation event. In addition, we recovered several double and triple mutants.

These results demonstrate that simultaneous deletion of four genes using CRISPR in *P. patens* is achievable in a single transformation. We next tested whether expressing gRNAs in a polycistronic construct could further enhance multiplex gene deletion efficiency. Although we previously found that polycistronic gRNA expression improves single-gene editing (e.g., *MAD2*), its effectiveness for multi-gene deletions remained uncertain.

We thus cloned the successful gRNA pairs from Construct 5–6 into a polycistronic construct (named Construct tRNA 5–6), where each gRNA was separated by tRNA spacers and driven by a single U6 promoter. PCR genotyping revealed that Construct tRNA 5–6 substantially improved editing efficiency: full deletions were detected in 32.5% of plants for *TPX2_1*, 34.5% for *TPX2_2*, 42.35% for *TPX2_3*, and 26.82% for *TPX2_4*. Using this optimized construct, we isolated nine quadruple mutants, as well as multiple triple and double mutants. Finally, we selected two quadruple mutants #6 and #9 to confirm genotyping PCR results by sequencing (Supplemental Figure 1, Supplemental dataset 2). As expected, large gene deletions were confirmed in all four TPX2 genes, with approximately ∼7780 bp deletion in *TPX2_1,* ∼6960 bp deletion in *TPX2_2,* 2266 bp deletion in *TPX2_3,* and ∼ 4710 bp deletion in *TPX2_4.*

These findings establish that polycistronic CRISPR constructs significantly enhance the efficiency of large, multiplex gene deletions in *P. patens*.

## Discussion

In this study, we evaluated CRISPR/Cas9-based strategies to induce large deletions in multiple genes simultaneously in *P. patens*. By employing dual-gRNA targeting, we successfully generated deletions encompassing the majority of the coding sequences of up to four distinct genes in a single transformation event. Traditionally, complete gene knockouts in *P. patens* have been achieved through highly efficient homologous recombination (Hohe et al., 2004; Kamisugi et al., 2006). This classical method typically involves replacing the target gene with an antibiotic resistance cassette.

However, homologous recombination is generally limited to removing a single gene per transformation and becomes increasingly impractical when targeting gene families with multiple paralogs. In such a case, the process is both time-consuming and labor-intensive, requiring sequential transformations and extensive molecular screening.

Moreover, the limited number of available antibiotic resistance cassettes poses an additional constraint when multiple rounds of transformation are needed. In contrast, CRISPR/Cas9-based approaches in *P. patens* typically rely on transient selection, which does not result in stable integration of the resistance cassette into the genome. This allows the same antibiotic to be reused in subsequent transformations of the obtained mutants.

While CRISPR-induced indels have been widely used for gene knockout in *P. patens* (Lopez-Obando et al., 2016; Trogu et al., 2021), full gene deletions spanning several kilobases have been reported less frequently and primarily in the context of individual loci. Nomura et al., 2016 reported deletions of up to 3 kb, Collonnier et al., 2017 achieved deletions of 3–4 kb, and more recently, Trogu et al., 2021 demonstrated deletions of up to 6 kb. Our study builds on this foundation by demonstrating the simultaneous induction of large deletions across four independent loci in a single transformation event. To our knowledge, this represents the first report in a plant system of coordinated, multi-locus deletions, ranging from 2 to 7 kb. These findings extend the capacity of CRISPR/Cas9 in *P. patens* and provide a scalable method for targeting gene families with high redundancy.

A key aspect of our approach was the implementation of a tRNA-based polycistronic gRNA expression system. This method, previously shown to enhance gRNA multiplexing efficiency in Arabidopsis and rice (Xie et al., 2015; Zhang et al., 2016), was adapted here for use in *P. patens*. Our results indicate that polycistronic constructs significantly improve editing efficiency compared to conventional designs where each gRNA is driven by a separate *PpU6* promoter. This improvement was consistent across both single and multi-gene editing contexts, suggesting that the tRNA system supports robust and coordinated expression of multiple gRNAs in moss.

On the other hand, we observed that deletion efficiency varied considerably between target loci and between different gRNA pairs. This variability likely reflects differences in gRNA cleavage efficiency, chromatin accessibility, or genomic context, consistent with known determinants of CRISPR activity (Isaac et al., 2016; Tycko et al., 2016).

This variability becomes increasingly problematic in multiplex designs, where the likelihood of at least one ineffective gRNA pair increases with the number of genes targeted.

We also evaluated whether increasing the number of gRNAs per gene improves deletion efficiency. While constructs with six gRNAs showed slightly higher frequencies of deletion, they also resulted in more partial deletions. This outcome may be due to variable activity across gRNAs or interference from overlapping cleavage events. Such findings align with previous concerns regarding excessive multiplexing, including elevated off-target risks and incomplete editing (Cameron et al., 2017). For practical purposes, we find that two high-quality gRNAs per gene, positioned at the 5′ and 3′ ends of the coding region, provide a suitable balance between simplicity and efficiency. Importantly, the efficiency of these gRNA pairs remains a limiting factor; several tested gRNAs failed to generate deletions entirely, emphasizing the need for empirical validation of candidate guides.

In summary, our results show that large, multiplexed gene deletions are achievable in *P. patens* using CRISPR/Cas9, and that efficiency is significantly improved by polycistronic gRNA expression. While further optimization is needed, particularly for gRNA efficiency prediction, this approach offers a versatile platform for generating high-order knockouts and will facilitate genetic studies in development, signaling, and synthetic biology applications in moss.

## Materials and Methods

### *P.patens* transformation and culture

*P. patens* was cultured on BCDAT medium under continuous white light at 25□°C. The BCDAT medium contained 1□mM MgSO□, 1.84□mM KH□PO□ (pH 6.5), 10□mM KNO□, 1□mM CaCl□, 5□mM ammonium tartrate, and a micronutrient solution (Alternative TES) comprising 0.22□μM CuSO□, 10□μM H□BO□, 0.23□μM CoCl□, 0.1□μM Na□MoO□, 0.19□μM ZnSO□, 2□μM MnCl□, and 0.17□μM KI, solidified with 0.8% (w/v) agar. For antibiotic selection hygromycin B (30□mg/L) was added as needed.

Protoplasts were prepared from 5–6-day-old protonemata by enzymatic digestion in 8% (w/v) mannitol with 2% driselase for 30□min at room temperature. Cells were filtered through a 50□μm nylon mesh and washed twice with 8% mannitol. Protoplasts were resuspended at 1.6□×□10□□cells/mL in MMM solution consisting of 450□mM mannitol, 15□mM MgCl□, and 5□mM MES (pH 5.6). For transformation, 30□μg/μl of circular plasmid DNA was added to 300□μL of protoplast suspension and gently mixed with 300□μL of PEG solution (8% PEG6000, 10□mM Ca(NO□)□, 10□mM Tris–HCl, pH 8.0). The mixture was incubated at 45□°C for 5□min followed by 10□min at 20□°C.

Transformation mixtures were gradually diluted with protoplast liquid medium composed of 66□mM mannitol, 27.8□mM glucose, 5.4□mM ammonium tartrate, 2.5□mM MgSO□, 0.184□mM KH□PO□, 0.5□mM Ca(NO□)□, and micronutrients (as above). After overnight incubation in darkness, protoplasts were harvested and resuspended in the warm PRM/T solution (400□mM mannitol, 5□mM ammonium tartrate, 1□mM MgSO□, 1.84□mM KH□PO□, 10□mM KNO□, 1□mM CaCl□, micronutrients, 0.8% low-melting agar). Protoplasts in PRM/T solution were then spread onto PRM plates (same as BCDAT medium with the 330□mM mannitol and sans ammonium tartrate) overlaid with sterile cellophane, and incubated for 2–4 days under continuous white light at 25□°C. Cellophanes were subsequently transferred to hygromycin-containing BCDAT medium for 5–7 days for antibiotic selection. Following selection, the cellophanes were moved to regular BCDAT plates and cultured until moss colonies regenerated to a suitable size for picking. Colonies were validated by PCR-based genotyping.

### *P.patens* genotyping

Transgenic colonies were genotyped by PCR to validate target gene deletion by CRISPR/Cas9. A small portion of moss colony (single gametophore or similar size of protonemata tissue) were picked from each colony and placed into individual wells of a 96-well PCR plate, each containing single plastic bead and 30□μL of 10×Green PCR buffer (670□mM Tris–HCl, 160□mM (NH□)□SO□, 0.1% Tween 20, pH 8.8). The plate was sealed, and moss tissues were homogenized using a tissue lyser (QIAGEN TissueLyser II; 1□min at 20□Hz), followed by centrifugation at 510□×□g for 15□min. The resulting supernatants were used directly as PCR templates.

PCR reactions (8□μL) were assembled using the KOD ONE polymerase (TOYOBO), with final concentrations of 4□μL 2× KOD One^®^ PCR Master Mix -Blue-, 4 μL distilled water, 0.125□μM forward and reverse primers, and 0.8□μL template. The PCR cycle parameters were: 35 cycles of denaturation at 98□°C for 10□s, primer annealing at 60-63□°C for 5□s, extension at 68□°C for 10-30 sec□per kb. PCR products were analyzed by agarose gel electrophoresis (3□μL PCR reaction per lane). Raw gel images are provided in the Supplemental dataset 1.

Primers used for genotyping of the *MAD2* gene (Pp3c4_13910V3.1; Pp6c4_7190V6.1), *katanin p60_1* (Pp3c15_10930V3.1; Pp6c15_5750V6.1), *katanin p60_2* (Pp3c9_3800V3.1; Pp6c9_2030V6.1), *TPX2_1* (Pp3c17_11160V3.1; Pp6c17_5570V6.1), *TPX2_2* (Pp3c1_25950V3.1; Pp6c1_13650V6.1), *TPX2_3* (Pp3c24_8590V3.1, Pp6c24_4210V6.3), *TPX2_4* (Pp3c23_4540V3.1; Pp6c23_1740V6.1) are provided in the Supplemental Table 1.

### gRNA design and cloning procedures

To generate CRISPR/Cas9-induced deletions in multiple genes, 20 bp guide RNAs (gRNAs) were designed to target the 5′ and 3′ coding regions of each gene using CRISPOR website (http://crispor.tefor.net/). Pairs of gRNAs were selected to induce large deletions spanning the majority of the coding sequence. Oligonucleotides bearing the gRNA sequences with appropriate overhangs were cloned into entry vectors using BsaI (for *MAD2*) or BbsI (for *katanin, TPX2*) restriction sites, depending on the modular system employed.

For constructs in which gRNAs were expressed individually, each gRNA was placed under the control of a *PpU6* promoter followed by the standard gRNA scaffold. For *MAD2* constructs, the region spanning the *PpU6* promoter to the end of the gRNA scaffold was PCR-amplified and assembled into a single vector carrying a hygromycin resistance cassette using Gibson Assembly. In these experiments, the Cas9 nuclease, driven by the rice *Actin1* promoter, was provided from a separate plasmid. Moss protoplasts were transformed with a 1:1 mixture of the gRNA-containing plasmid and the Cas9 expression vector.

Polycistronic *MAD2* constructs were generated using Golden Gate Assembly. Each gRNA-gRNA scaffold was flanked by tRNA sequences. PCR fragments were amplified from the EKv376 template, and Golden Gate primers were designed to amplify approximately half of each gRNA followed by BsaI recognition sites. This strategy allowed the complete gRNA sequences to be reconstituted upon BsaI digestion and ligation during the Golgen Gate reaction.

Notably, we found that assembling polycistronic constructs using in-house Golden Gate reactions was technically challenging and time-consuming. Therefore, for the *TPX2* polycistronic gRNA vector, we opted for commercial gene synthesis (GENEWIZ from Azenta Life Sciences) instead of in-house cloning.

### Modular CRISPR/Cas9 system for Golden Gate assembly

We designed a vector system that utilizes Golden Gate assembly for efficient and scalable construction of multiplex gRNA arrays. Our system supports the one-step assembly of 4, 6, or 8 gRNA expression cassettes into a single binary vector.

Each gRNA was initially cloned into an intermediate vector, called gRNA insert vectors, by annealing complementary oligonucleotides with appropriate overhangs and inserting them via BbsI-HF(New England Biolabs) digestion into a site located between a *PpU6* promoter and the gRNA scaffold. The *PpU6* promoter and gRNA scaffold were flanked by outward-facing BsaI recognition sites with unique four-base overhangs to direct ordered assembly. Once verified by Sanger sequencing, the individual gRNA vectors were used for Golden Gate-based assembly into a destination vector.

The destination vector was designed to accept multiple gRNA units through BsaI digestion and ligation. It also included a Cas9 nuclease driven by the *lhcsr1* promoter (Hiss et al., 2017), in which all internal BsaI sites were removed by silent mutations to preserve compatibility with Golden Gate assembly. A hygromycin resistance cassette under the control of the CaMV 35S promoter was included to allow selection in moss.

Golden Gate assembly was performed in a 15□μL reaction containing 1□μL of the destination vector (e.g., EKv444), 1□μL of each gRNA insert vector, 1.5□μL of 10× T4 DNA ligase buffer, 1.5□μL of BsaI-HF (New England Biolabs), and 0.5□μL of T4 DNA ligase (New England Biolabs). Nuclease-free water was added to bring the total volume to 15□μL. Reaction components were added in the following order: destination vector and gRNA inserts, followed by BsaI-HF and ligase buffer, and finally T4 DNA ligase. The reaction was incubated in a thermocycler using the following program: 37□°C for 30□min; followed by 35 cycles of 37□°C for 5□min and 16□°C for 5□min; then 37□°C for 30□min, 60□°C for 5□min, and held at 8□°C. Assembled products were transformed into *E. coli* and screened by PmeI (New England Biolabs) digestion, followed by sequencing across the gRNA array region. Correctly assembled multiplex CRISPR/Cas9 constructs were used for PEG-mediated transformation of *P. patens* protoplasts.

## Conflict of interest

The authors declare no competing interests.

## Contributions

E.K. designed and executed this research project, acquired funding, wrote and edited the manuscript.

## Acknowledgements

We would like to thank Dr. Moé Yamada for providing GPH1029 *P. patens* line, Rie Inaba for performing *P. patens* genetic transformation and genotyping PCR. This work was funded by Japan Society for the Promotion of Science (KAKENHI) 23K14210 and Research Grant for Young Japanese Researchers from the Nakajima Foundation to EK.

---

**Supplemental Figure 1.**
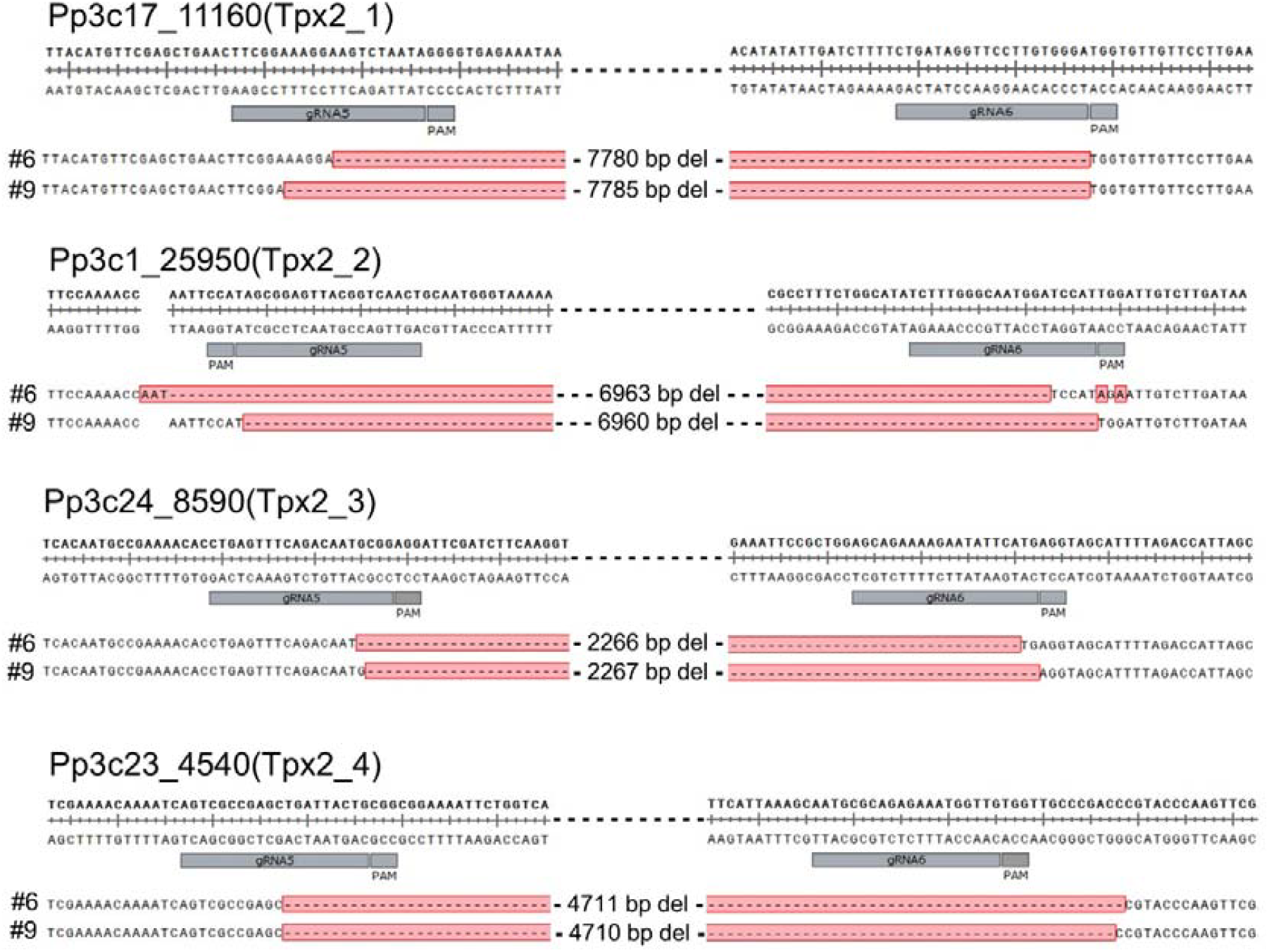
Sequencing confirms large deletions in coding regions of four TPX2 genes. Quadruple mutants #6 and #9 obtained after P. patens transformation with Construct tRNA 5-6 were selected for sequencing. The regions around 5’ and 3’ gRNA target sites are shown. Actual sequencing data is available in Supplemental Dataset 2.

**Supplemental Table 1**. gRNAs, primers, plasmids, and moss lines used in this study.

**Supplemental Dataset 1.** All genotyping PCR results from this study.

**Supplemental Dataset 2.** Sequencing data of TPX2 quadruple mutants #6 and #9.

## Notes

### Competing Interest Statement

The authors have declared no competing interest.

