## Supplemental dataset 1 for "tRNA-Based Polycistronic CRISPR/Cas9 System Boosts Efficiency of Multi-Gene Deletion in the Moss Physcomitrella": Supplemental dataset 1.pptx

### Slide 1
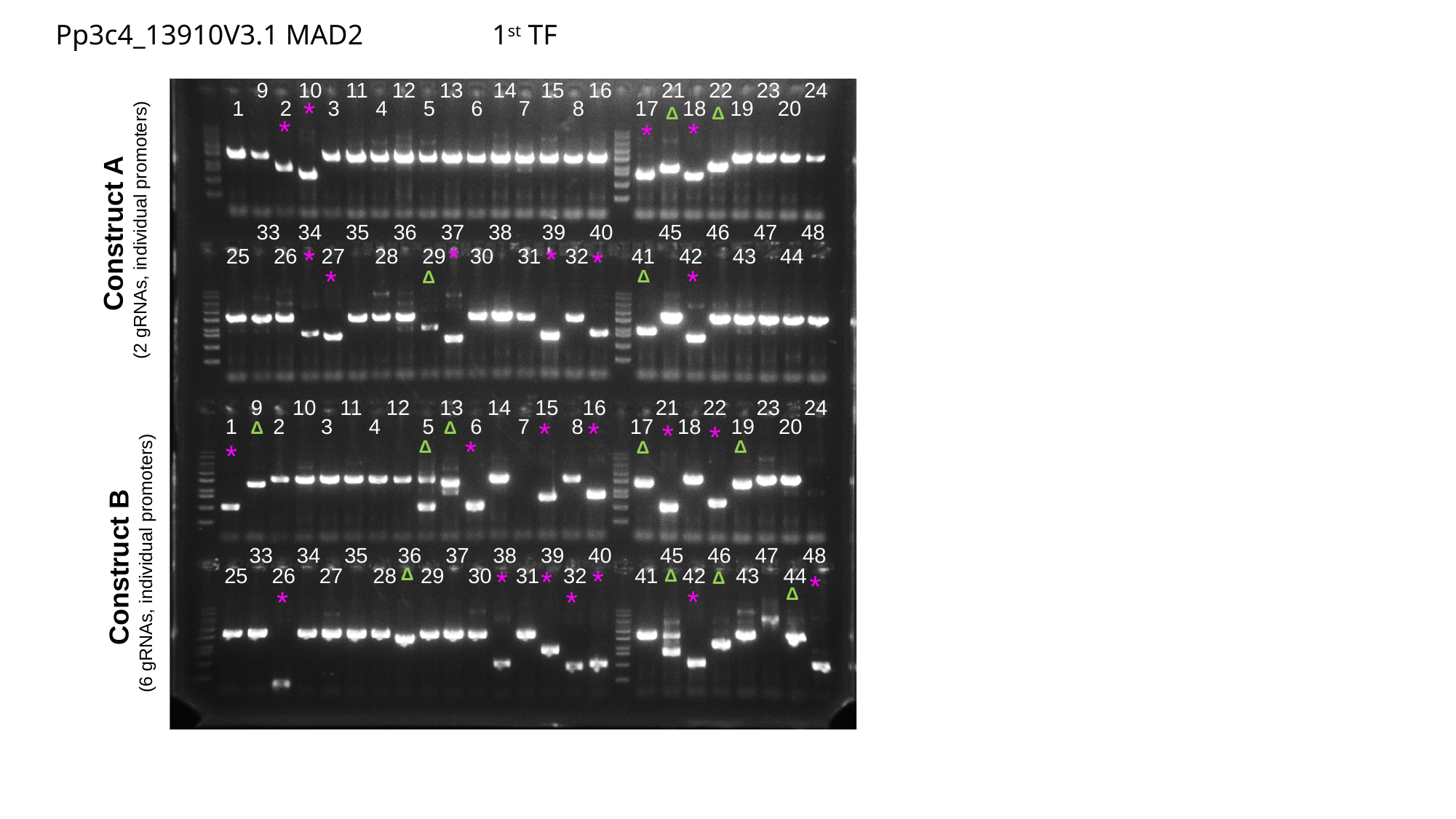

Pp3c4_13910V3.1 MAD2 		1^st^ TF
9 10 11 12 13 14 15 16
21 22 23 24
*
1 2 3 4 5 6 7 8
17 18 19 20
Δ
Δ
*
*
*
TF884
Construct A
(2 gRNAs, individual promoters)
33 34 35 36 37 38 39 40
45 46 47 48
*
*
*
*
25 26 27 28 29 30 31 32
41 42 43 44
*
*
Δ
Δ
9 10 11 12 13 14 15 16
21 22 23 24
1 2 3 4 5 6 7 8
17 18 19 20
*
*
Δ
Δ
*
*
*
Δ
Δ
Δ
*
Construct B
(6 gRNAs, individual promoters)
33 34 35 36 37 38 39 40
45 46 47 48
*
25 26 27 28 29 30 31 32
Δ
41 42 43 44
*
*
Δ
Δ
*
Δ
*
*
*

### Slide 2
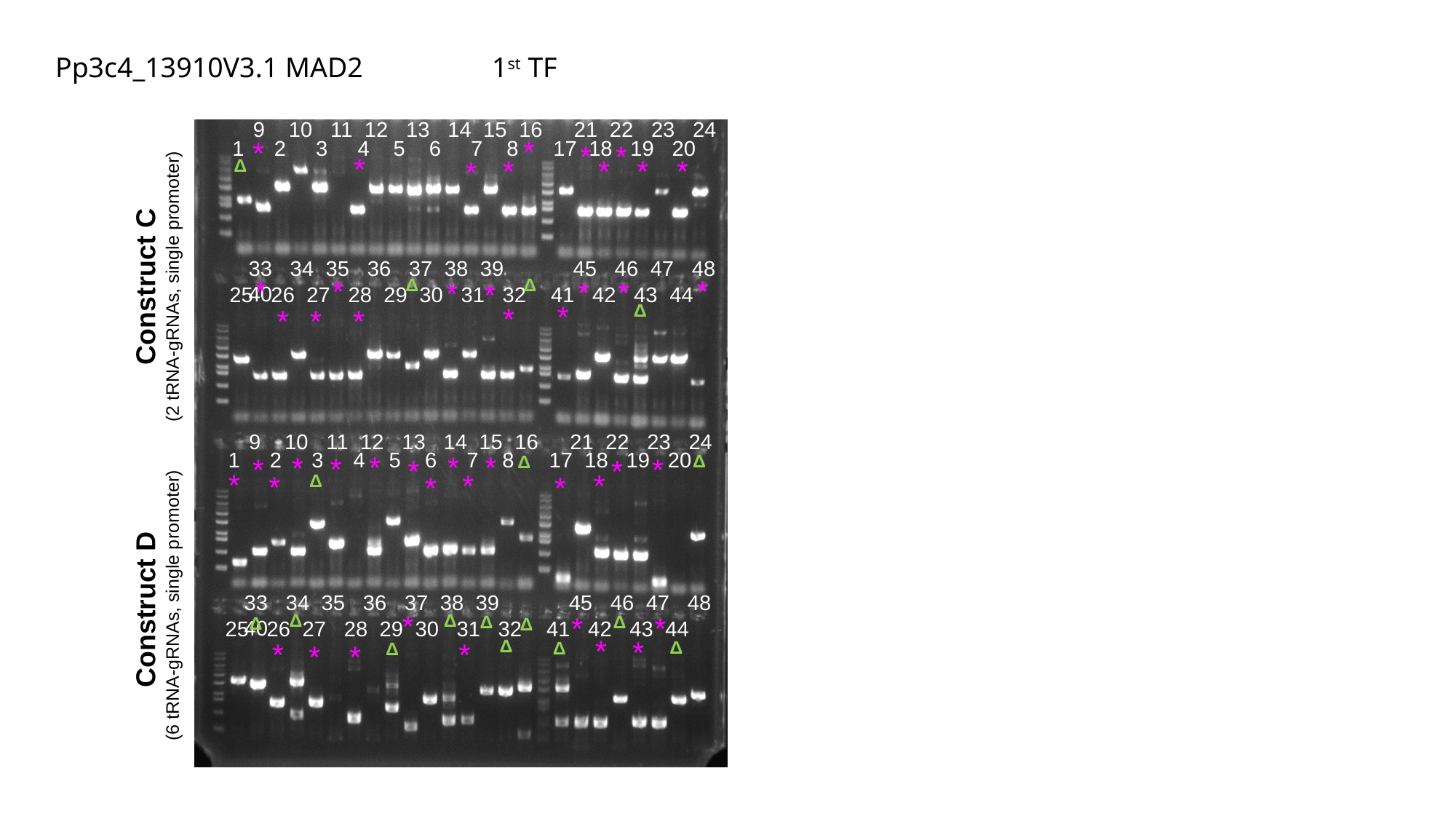

Pp3c4_13910V3.1 MAD2 		1^st^ TF
9 10 11 12 13 14 15 16
21 22 23 24
*
*
1 2 3 4 5 6 7 8
17 18 19 20
*
*
*
*
*
*
*
*
Δ
33 34 35 36 37 38 39 40
45 46 47 48
Construct C
(2 tRNA-gRNAs, single promoter)
*
*
*
*
*
Δ
Δ
*
*
25 26 27 28 29 30 31 32
41 42 43 44
*
Δ
*
*
*
*
9 10 11 12 13 14 15 16
21 22 23 24
1 2 3 4 5 6 7 8
17 18 19 20
*
*
*
*
*
Δ
Δ
*
*
*
*
*
*
*
*
*
*
Δ
Construct D
(6 tRNA-gRNAs, single promoter)
33 34 35 36 37 38 39 40
45 46 47 48
*
*
Δ
Δ
*
Δ
Δ
Δ
Δ
25 26 27 28 29 30 31 32
41 42 43 44
*
*
Δ
*
*
Δ
Δ
*
Δ
*

### Slide 3
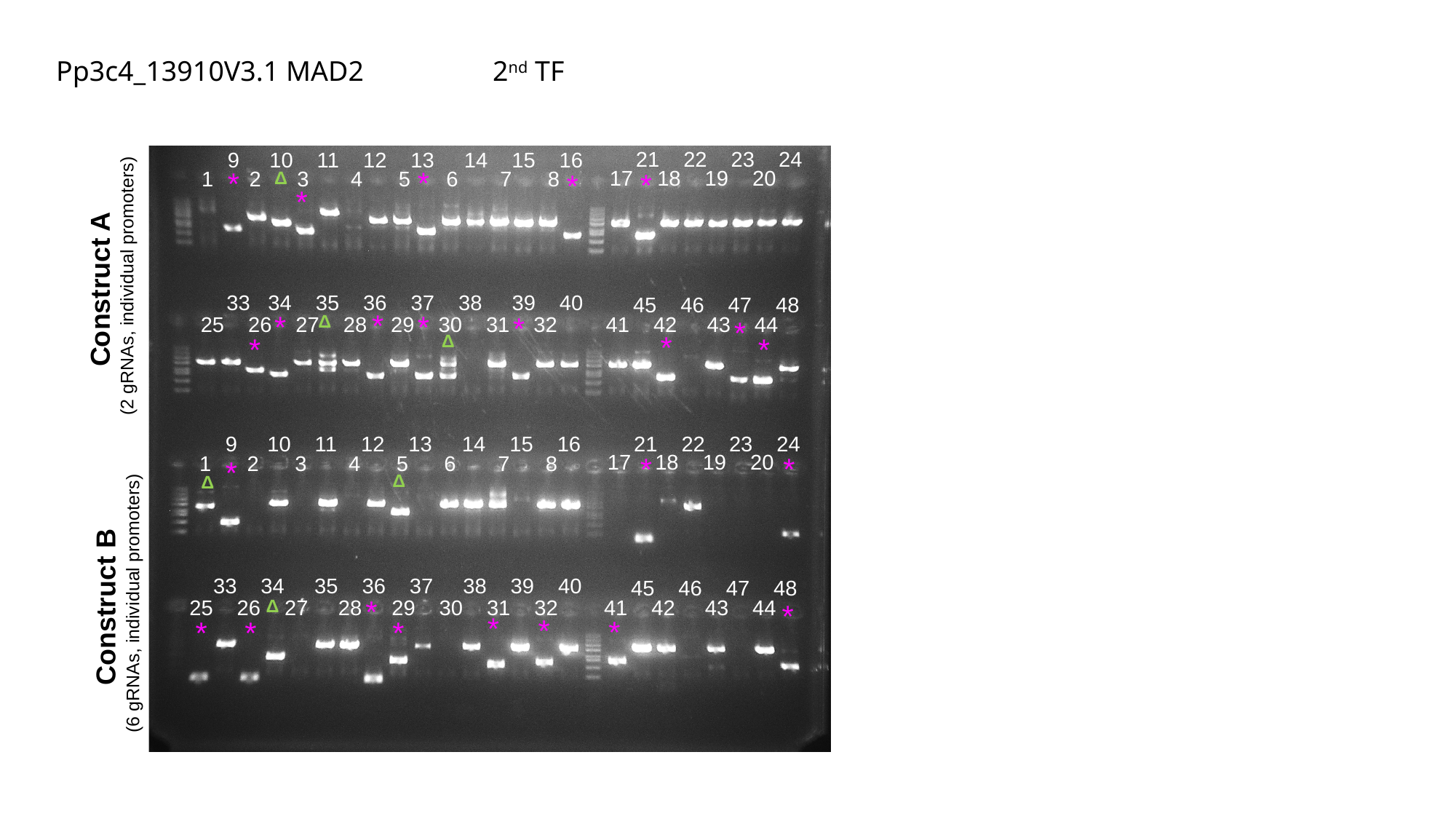

Pp3c4_13910V3.1 MAD2 		2^nd^ TF
21 22 23 24
9 10 11 12 13 14 15 16
*
*
17 18 19 20
1 2 3 4 5 6 7 8
*
Δ
*
*
Construct A
(2 gRNAs, individual promoters)
33 34 35 36 37 38 39 40
45 46 47 48
*
*
*
*
Δ
25 26 27 28 29 30 31 32
41 42 43 44
*
*
Δ
*
*
21 22 23 24
9 10 11 12 13 14 15 16
17 18 19 20
*
*
1 2 3 4 5 6 7 8
*
Δ
Δ
33 34 35 36 37 38 39 40
45 46 47 48
Construct B
(6 gRNAs, individual promoters)
*
25 26 27 28 29 30 31 32
41 42 43 44
Δ
*
*
*
*
*
*
*

### Slide 4
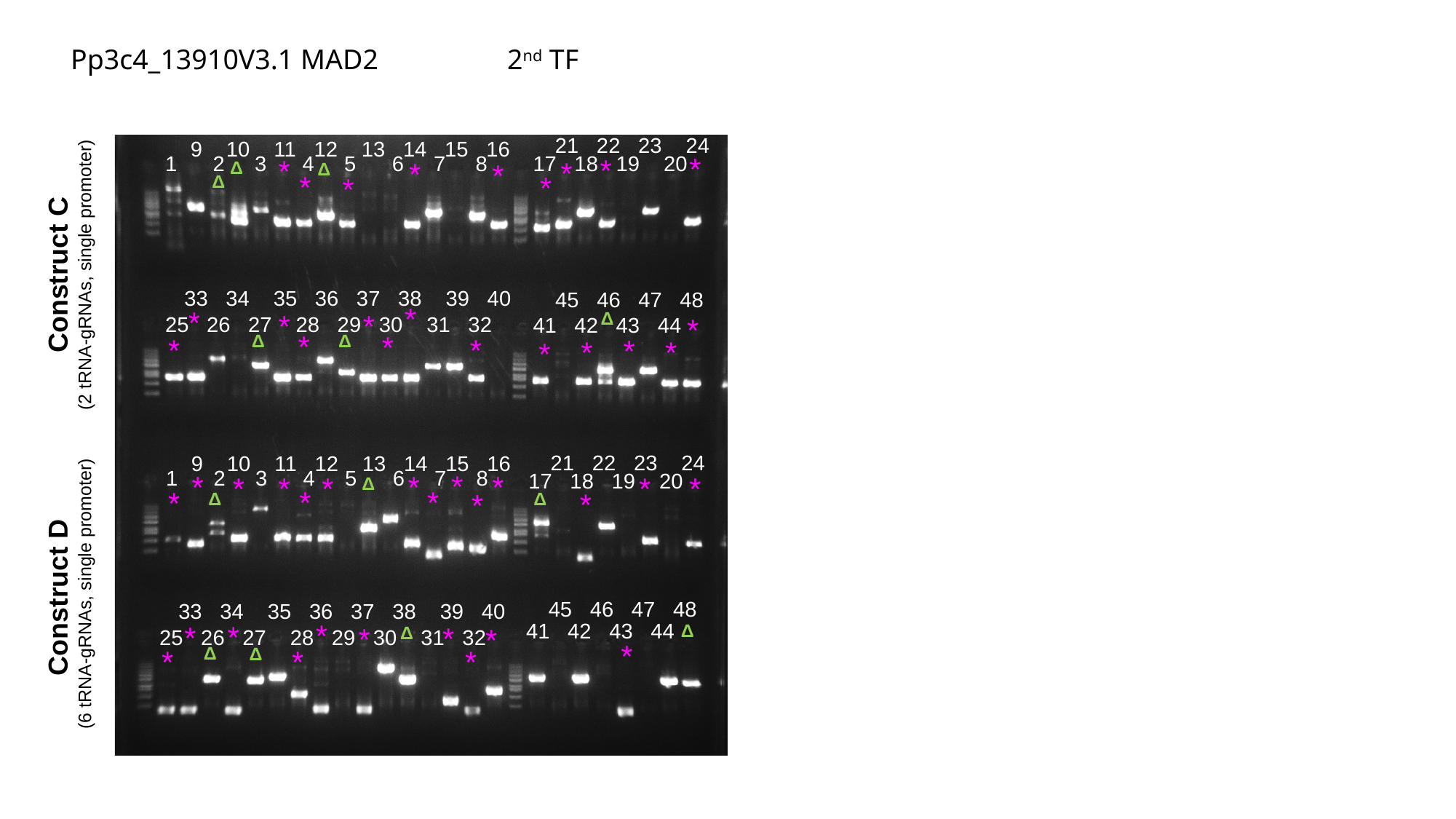

Pp3c4_13910V3.1 MAD2 		2^nd^ TF
21 22 23 24
9 10 11 12 13 14 15 16
*
1 2 3 4 5 6 7 8
17 18 19 20
*
*
*
*
Δ
*
Δ
*
*
Δ
*
Construct C
(2 tRNA-gRNAs, single promoter)
33 34 35 36 37 38 39 40
45 46 47 48
*
*
Δ
*
*
*
25 26 27 28 29 30 31 32
41 42 43 44
*
*
Δ
Δ
*
*
*
*
*
*
21 22 23 24
9 10 11 12 13 14 15 16
1 2 3 4 5 6 7 8
*
17 18 19 20
*
*
*
*
*
*
*
*
Δ
*
*
*
*
*
Δ
Δ
Construct D
(6 tRNA-gRNAs, single promoter)
45 46 47 48
33 34 35 36 37 38 39 40
*
*
41 42 43 44
*
Δ
*
*
*
Δ
25 26 27 28 29 30 31 32
*
Δ
Δ
*
*
*

### Slide 5
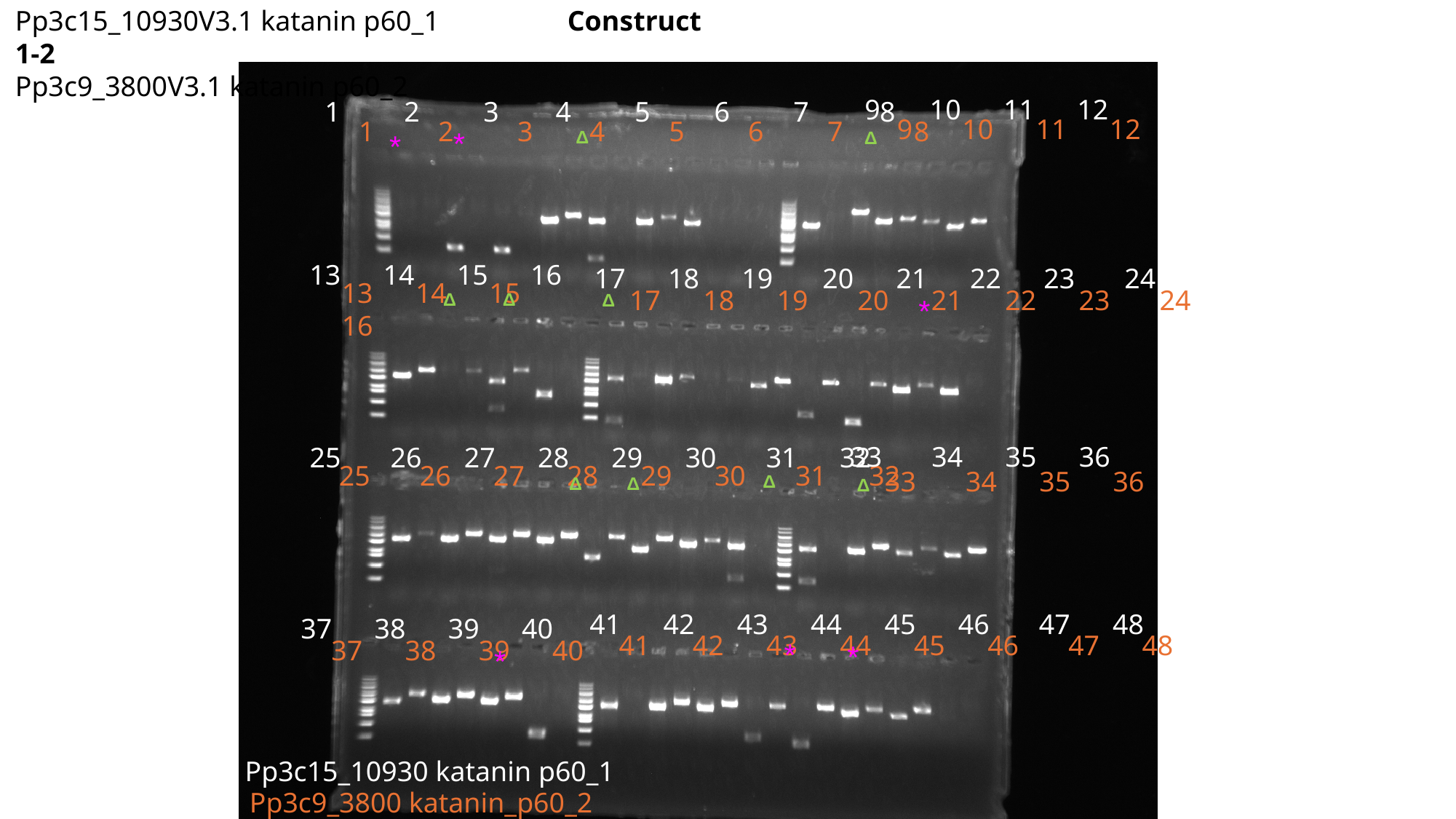

Pp3c15_10930V3.1 katanin p60_1 Construct 1-2
Pp3c9_3800V3.1 katanin p60_2
 9 10 11 12
1 2 3 4 5 6 7 8
 9 10 11 12
1 2 3 4 5 6 7 8
*
Δ
Δ
*
13 14 15 16
17 18 19 20 21 22 23 24
13 14 15 16
17 18 19 20 21 22 23 24
Δ
Δ
Δ
*
33 34 35 36
25 26 27 28 29 30 31 32
25 26 27 28 29 30 31 32
33 34 35 36
Δ
Δ
Δ
Δ
41 42 43 44 45 46 47 48
37 38 39 40
41 42 43 44 45 46 47 48
37 38 39 40
*
*
*
Pp3c15_10930 katanin p60_1
Pp3c9_3800 katanin_p60_2

### Slide 6
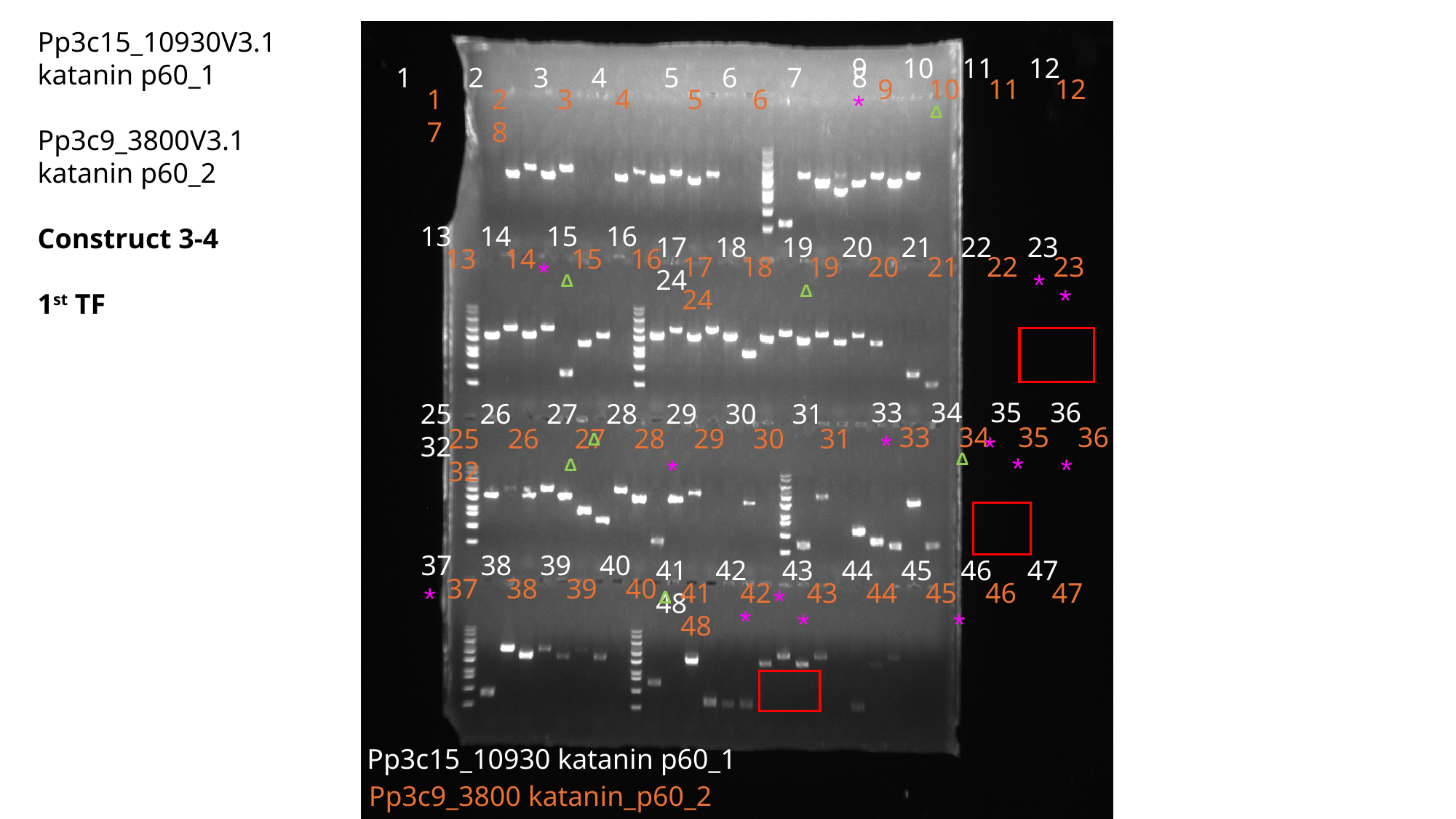

Pp3c15_10930V3.1 katanin p60_1 Pp3c9_3800V3.1 katanin p60_2
Construct 3-4
1^st^ TF
9 10 11 12
1 2 3 4 5 6 7 8
9 10 11 12
1 2 3 4 5 6 7 8
*
Δ
13 14 15 16
17 18 19 20 21 22 23 24
13 14 15 16
17 18 19 20 21 22 23 24
*
*
Δ
Δ
*
33 34 35 36
25 26 27 28 29 30 31 32
33 34 35 36
25 26 27 28 29 30 31 32
*
Δ
*
Δ
*
*
*
Δ
37 38 39 40
41 42 43 44 45 46 47 48
37 38 39 40
41 42 43 44 45 46 47 48
*
*
Δ
*
*
*
Pp3c15_10930 katanin p60_1
Pp3c9_3800 katanin_p60_2

### Slide 7
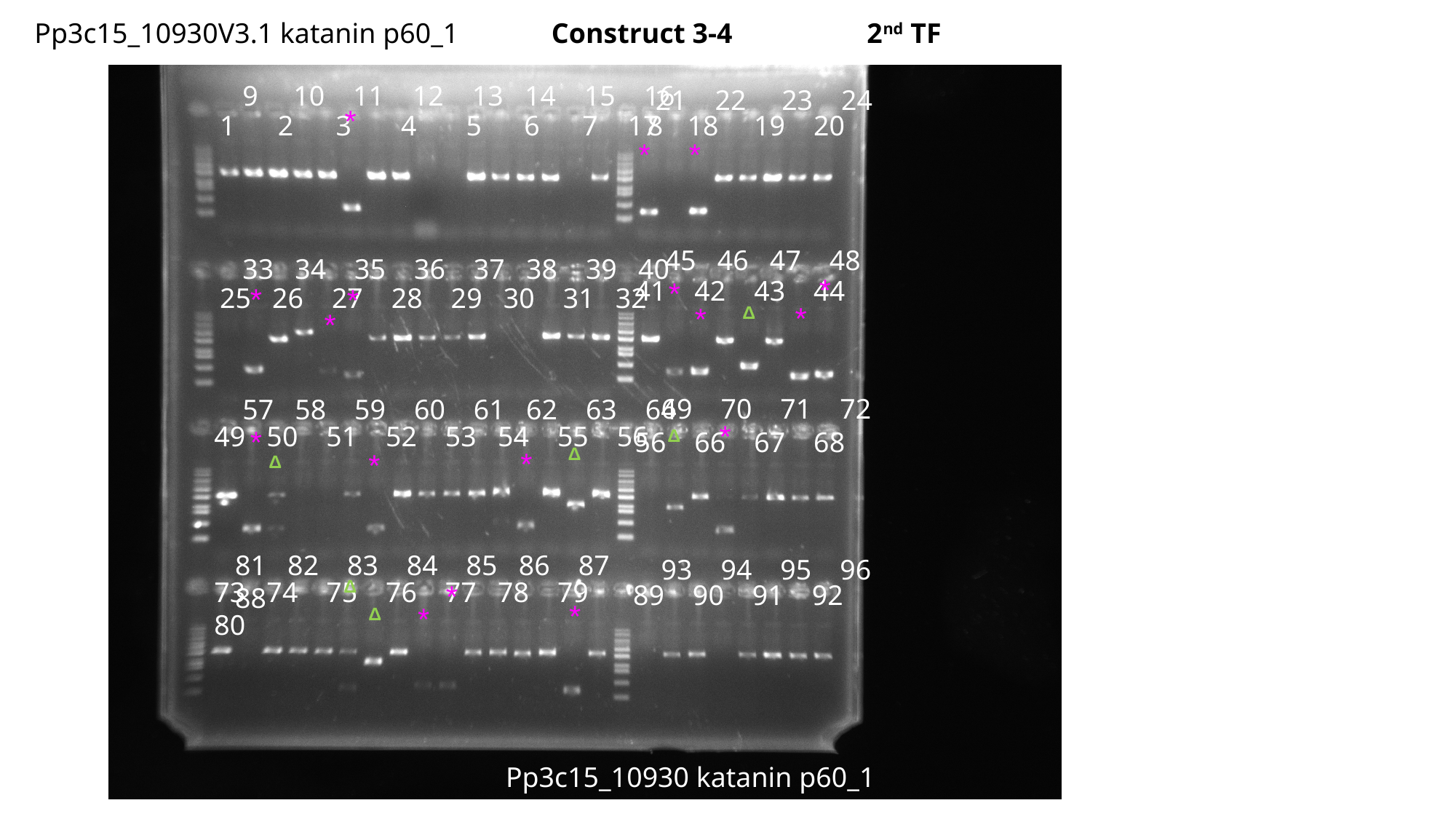

Pp3c15_10930V3.1 katanin p60_1 Construct 3-4 2^nd^ TF
9 10 11 12 13 14 15 16
21 22 23 24
*
1 2 3 4 5 6 7 8
17 18 19 20
*
*
45 46 47 48
33 34 35 36 37 38 39 40
*
41 42 43 44
*
*
*
25 26 27 28 29 30 31 32
*
*
Δ
*
69 70 71 72
57 58 59 60 61 62 63 64
*
49 50 51 52 53 54 55 56
*
Δ
56 66 67 68
Δ
*
*
Δ
81 82 83 84 85 86 87 88
93 94 95 96
Δ
73 74 75 76 77 78 79 80
*
89 90 91 92
*
*
Δ
Pp3c15_10930 katanin p60_1

### Slide 8
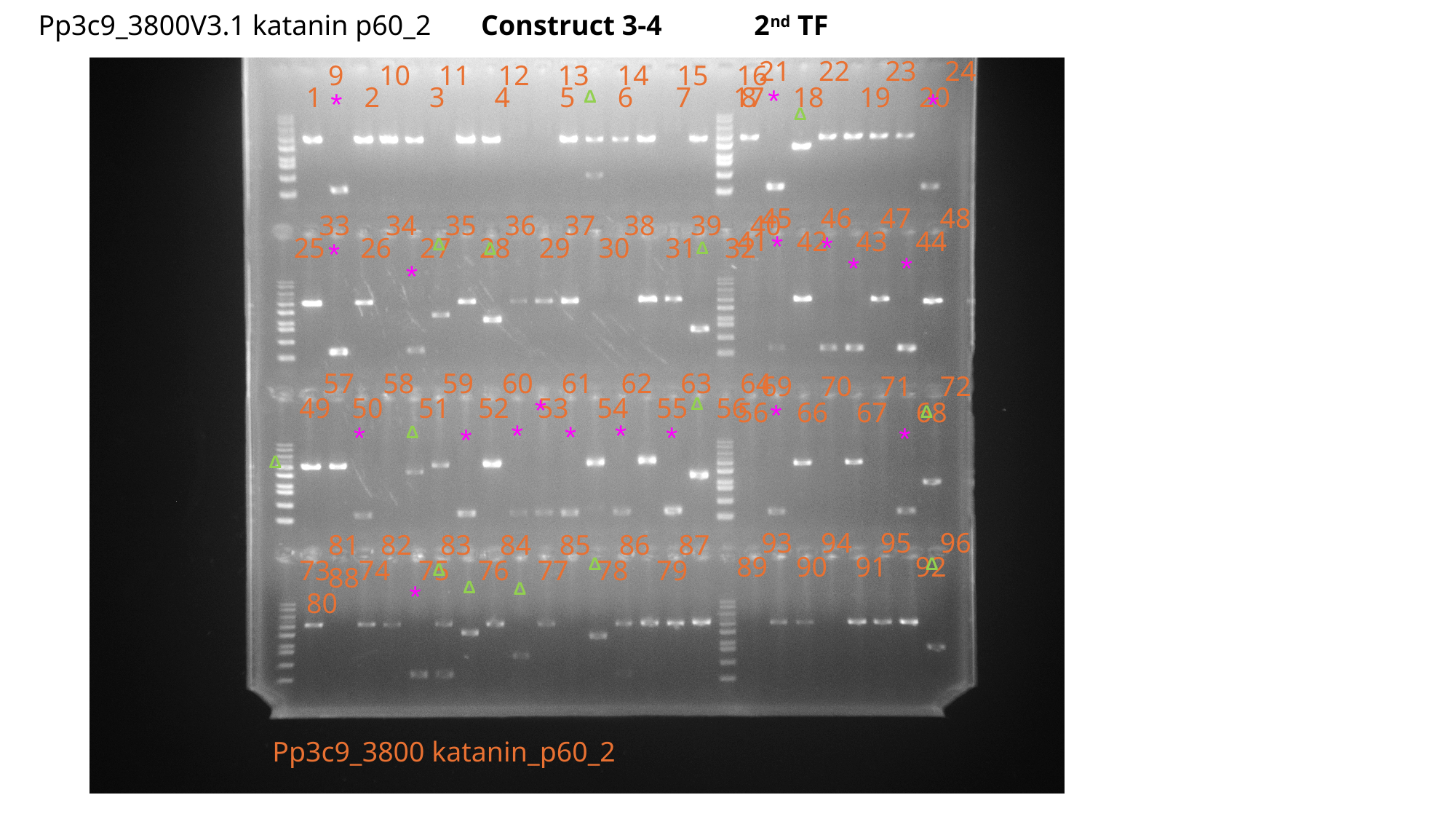

Pp3c9_3800V3.1 katanin p60_2 Construct 3-4 2^nd^ TF
21 22 23 24
9 10 11 12 13 14 15 16
1 2 3 4 5 6 7 8
17 18 19 20
*
Δ
*
*
Δ
45 46 47 48
33 34 35 36 37 38 39 40
41 42 43 44
*
*
25 26 27 28 29 30 31 32
Δ
*
Δ
Δ
*
*
*
57 58 59 60 61 62 63 64
69 70 71 72
*
49 50 51 52 53 54 55 56
Δ
56 66 67 68
*
Δ
*
*
*
*
*
*
Δ
*
Δ
93 94 95 96
81 82 83 84 85 86 87 88
89 90 91 92
Δ
Δ
73 74 75 76 77 78 79 80
Δ
Δ
Δ
*
Pp3c9_3800 katanin_p60_2

### Slide 9
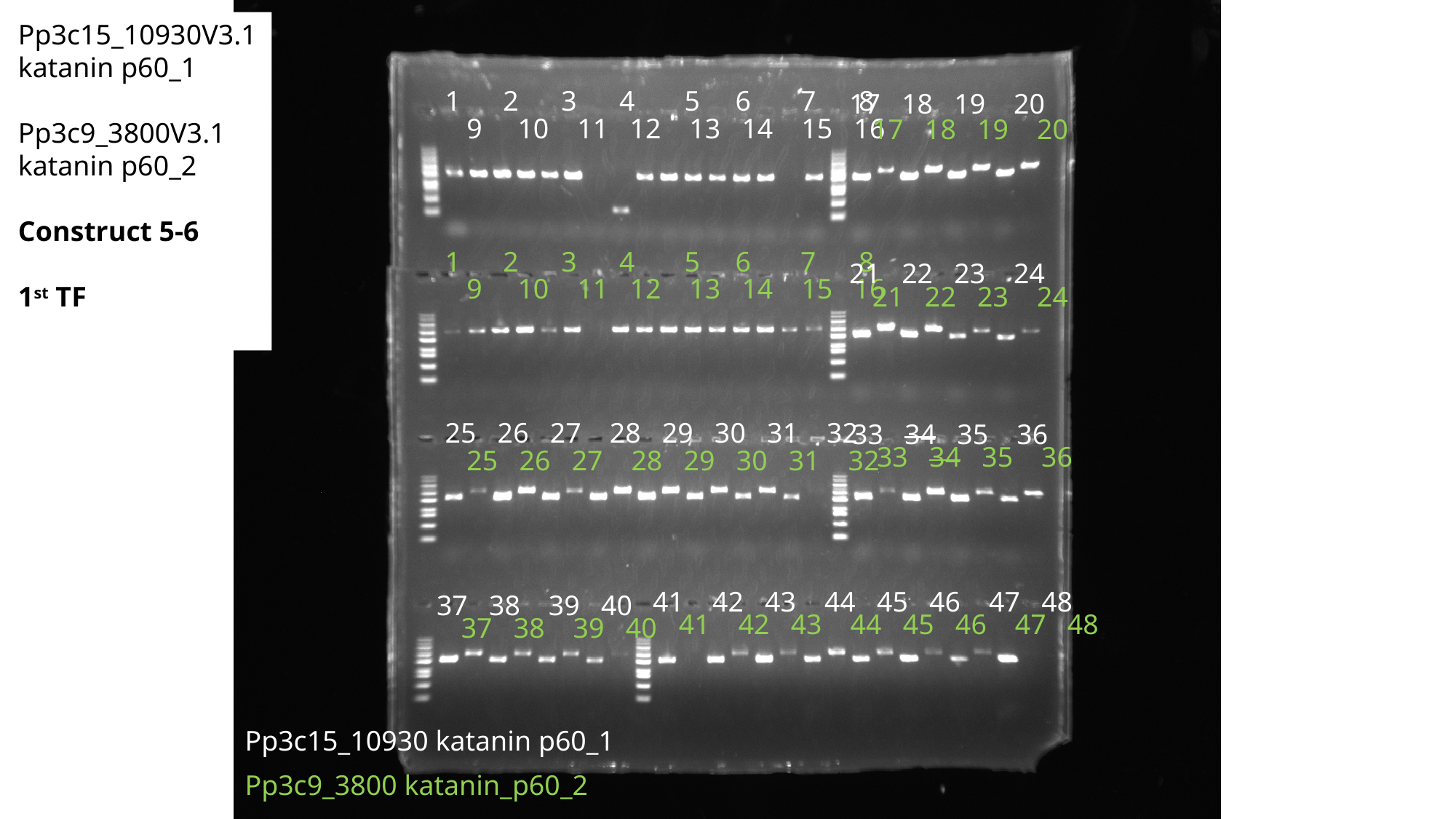

Pp3c15_10930V3.1 katanin p60_1 Pp3c9_3800V3.1 katanin p60_2
Construct 5-6
1^st^ TF
1 2 3 4 5 6 7 8
17 18 19 20
9 10 11 12 13 14 15 16
17 18 19 20
1 2 3 4 5 6 7 8
21 22 23 24
9 10 11 12 13 14 15 16
21 22 23 24
25 26 27 28 29 30 31 32
33 34 35 36
33 34 35 36
25 26 27 28 29 30 31 32
41 42 43 44 45 46 47 48
37 38 39 40
41 42 43 44 45 46 47 48
37 38 39 40
Pp3c15_10930 katanin p60_1
Pp3c9_3800 katanin_p60_2

### Slide 10
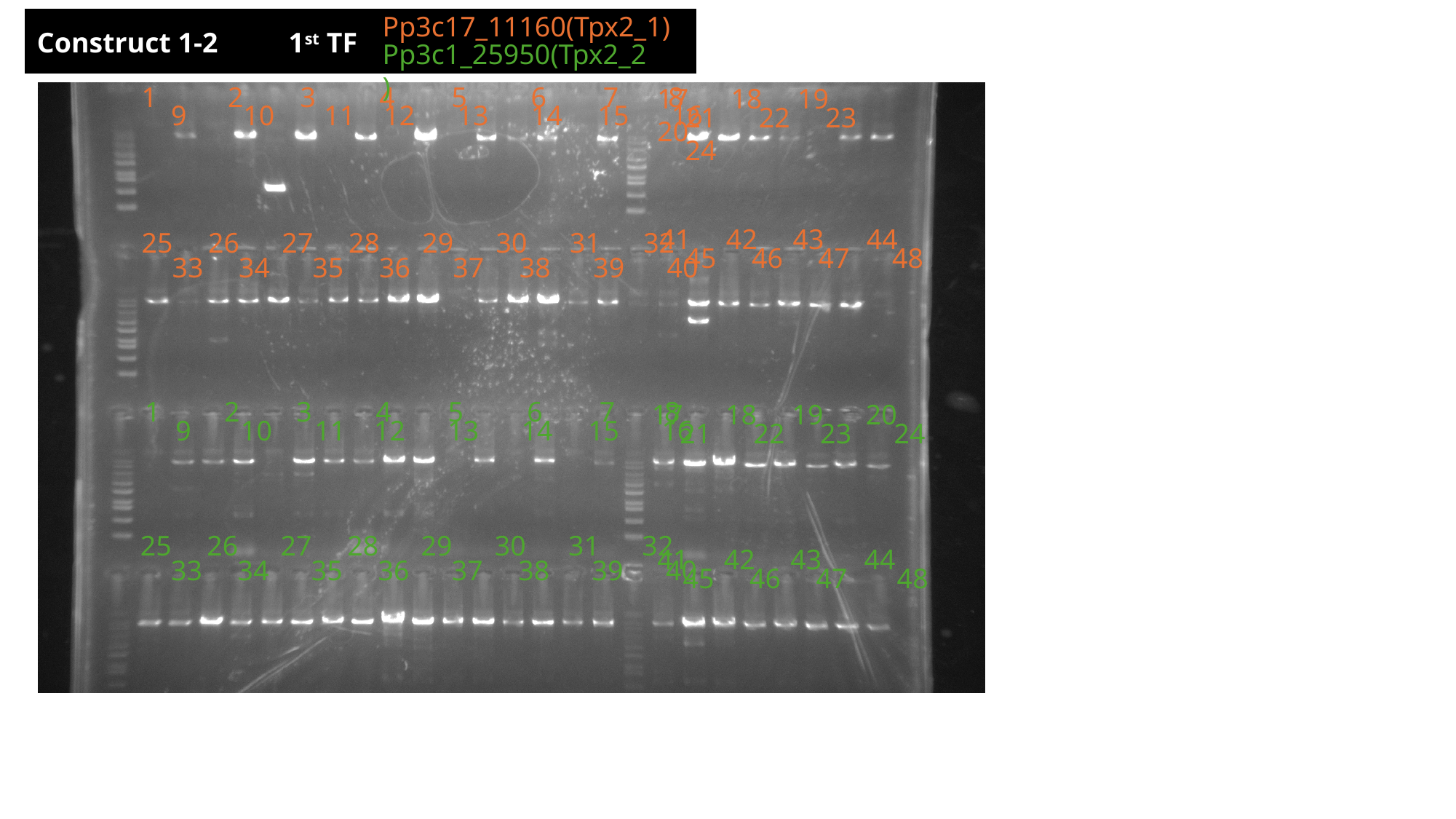

Pp3c17_11160(Tpx2_1)
Construct 1-2 1^st^ TF
Pp3c1_25950(Tpx2_2)
1 2 3 4 5 6 7 8
17 18 19 20
9 10 11 12 13 14 15 16
21 22 23 24
41 42 43 44
25 26 27 28 29 30 31 32
45 46 47 48
33 34 35 36 37 38 39 40
1 2 3 4 5 6 7 8
17 18 19 20
9 10 11 12 13 14 15 16
21 22 23 24
25 26 27 28 29 30 31 32
41 42 43 44
33 34 35 36 37 38 39 40
45 46 47 48

### Slide 11
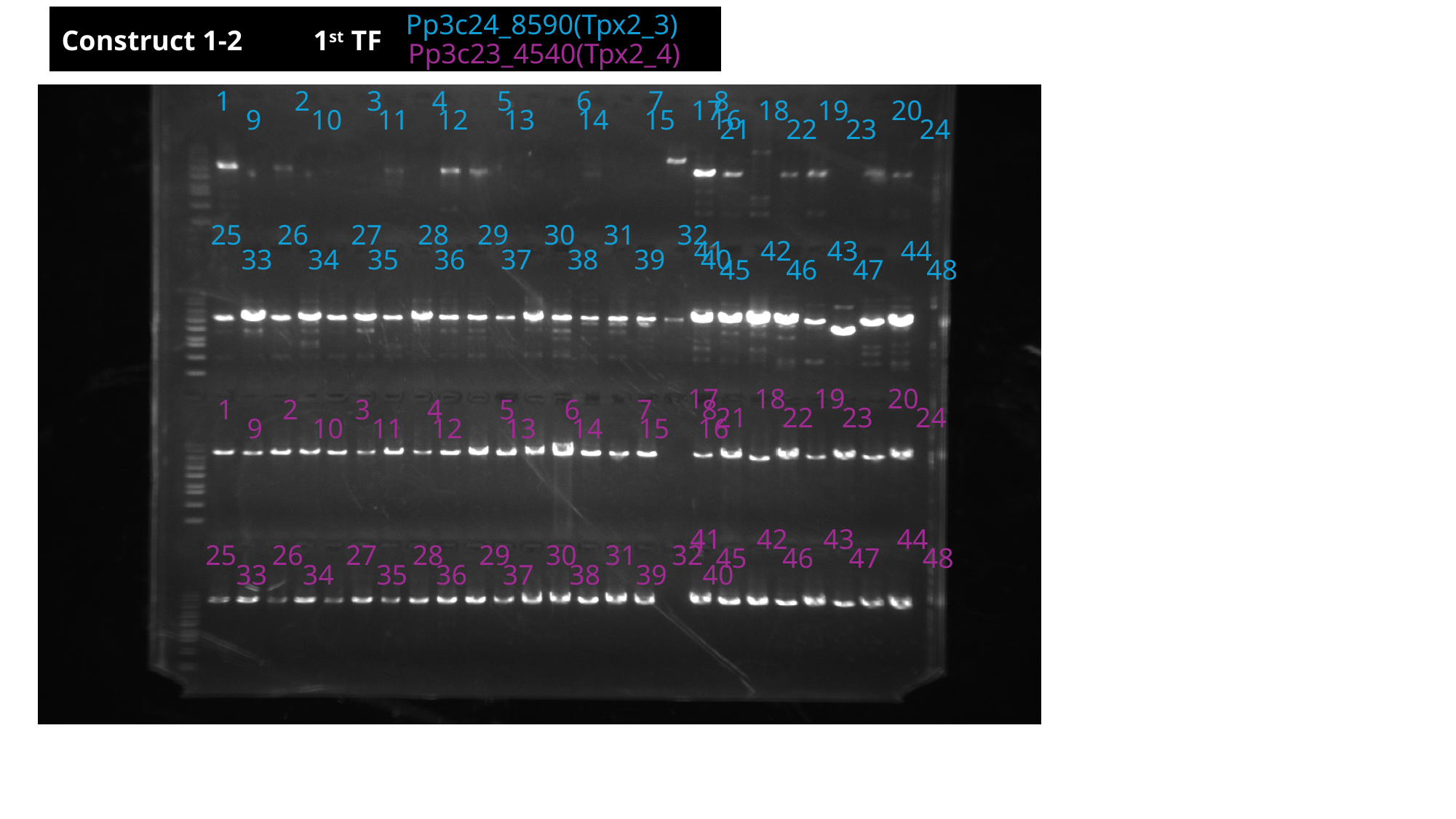

Pp3c24_8590(Tpx2_3)
Construct 1-2 1^st^ TF
Pp3c23_4540(Tpx2_4)
1 2 3 4 5 6 7 8
17 18 19 20
9 10 11 12 13 14 15 16
21 22 23 24
25 26 27 28 29 30 31 32
41 42 43 44
33 34 35 36 37 38 39 40
45 46 47 48
17 18 19 20
1 2 3 4 5 6 7 8
21 22 23 24
9 10 11 12 13 14 15 16
41 42 43 44
25 26 27 28 29 30 31 32
45 46 47 48
33 34 35 36 37 38 39 40

### Slide 12
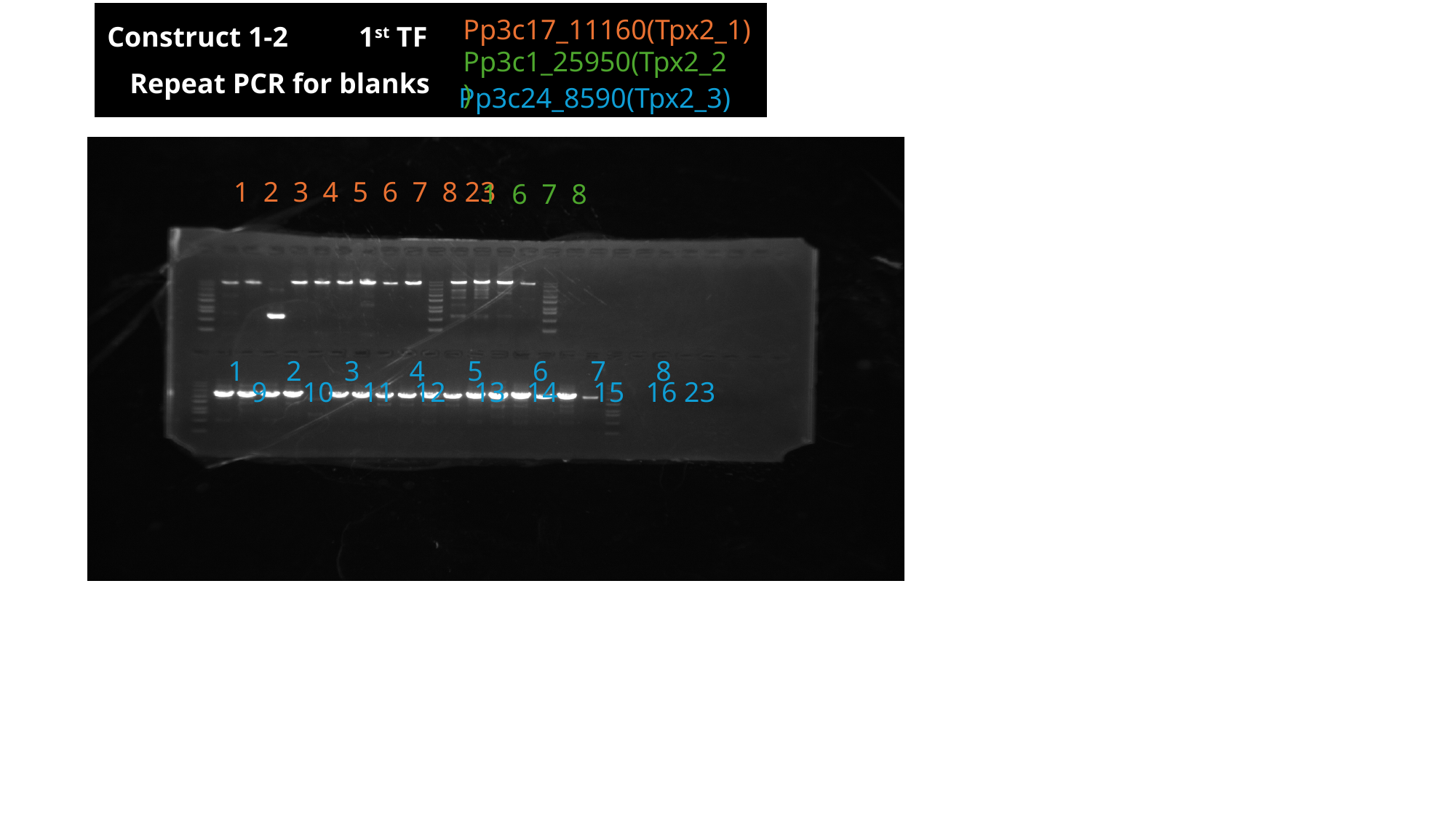

Pp3c17_11160(Tpx2_1)
Construct 1-2 1^st^ TF
Pp3c1_25950(Tpx2_2)
Repeat PCR for blanks
Pp3c24_8590(Tpx2_3)
 1 2 3 4 5 6 7 8 23
 1 6 7 8
1 2 3 4 5 6 7 8
9 10 11 12 13 14 15 16 23

### Slide 13
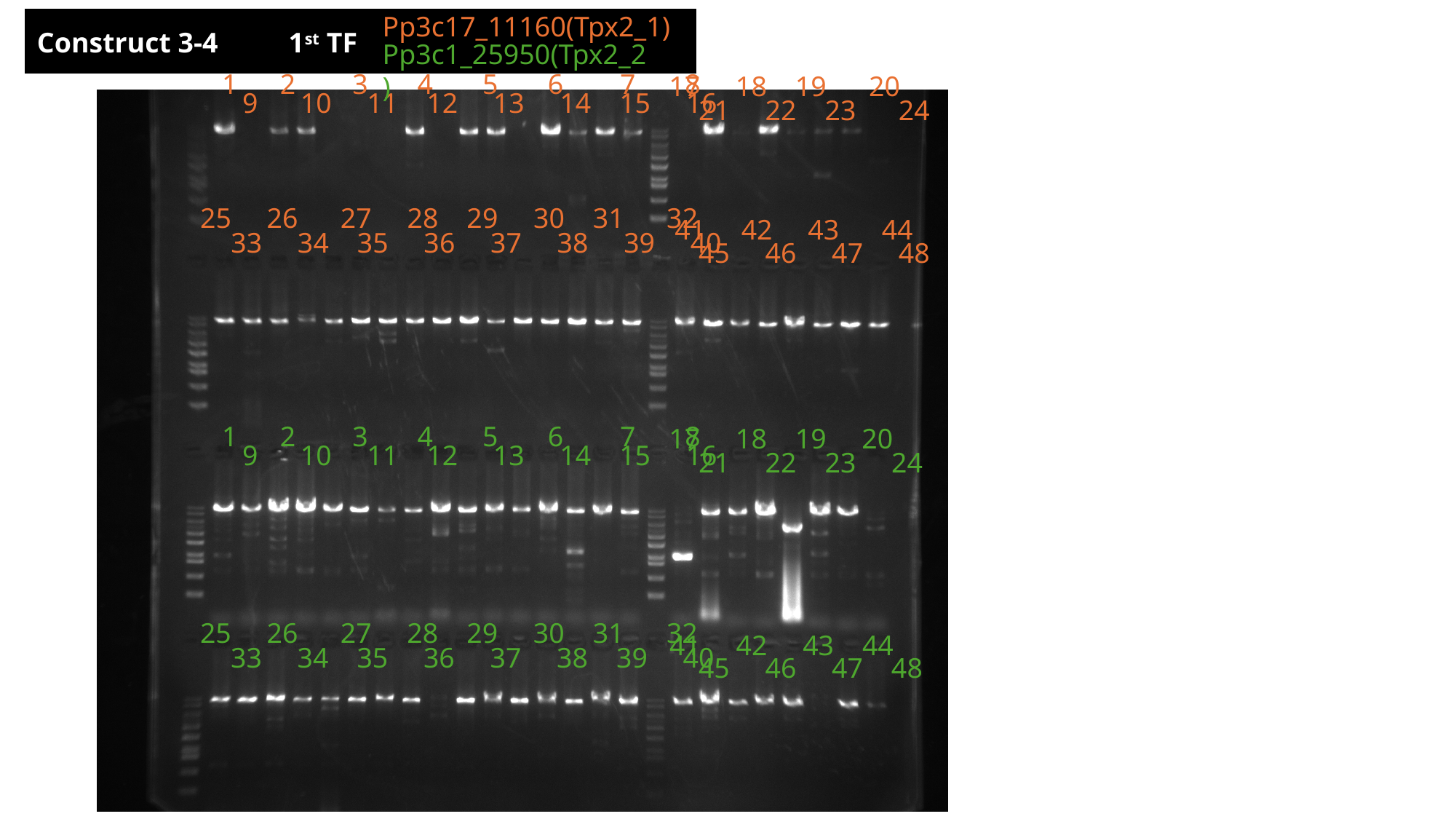

Pp3c17_11160(Tpx2_1)
Construct 3-4 1^st^ TF
Pp3c1_25950(Tpx2_2)
1 2 3 4 5 6 7 8
17 18 19 20
9 10 11 12 13 14 15 16
21 22 23 24
25 26 27 28 29 30 31 32
41 42 43 44
33 34 35 36 37 38 39 40
45 46 47 48
1 2 3 4 5 6 7 8
17 18 19 20
9 10 11 12 13 14 15 16
21 22 23 24
25 26 27 28 29 30 31 32
41 42 43 44
33 34 35 36 37 38 39 40
45 46 47 48

### Slide 14
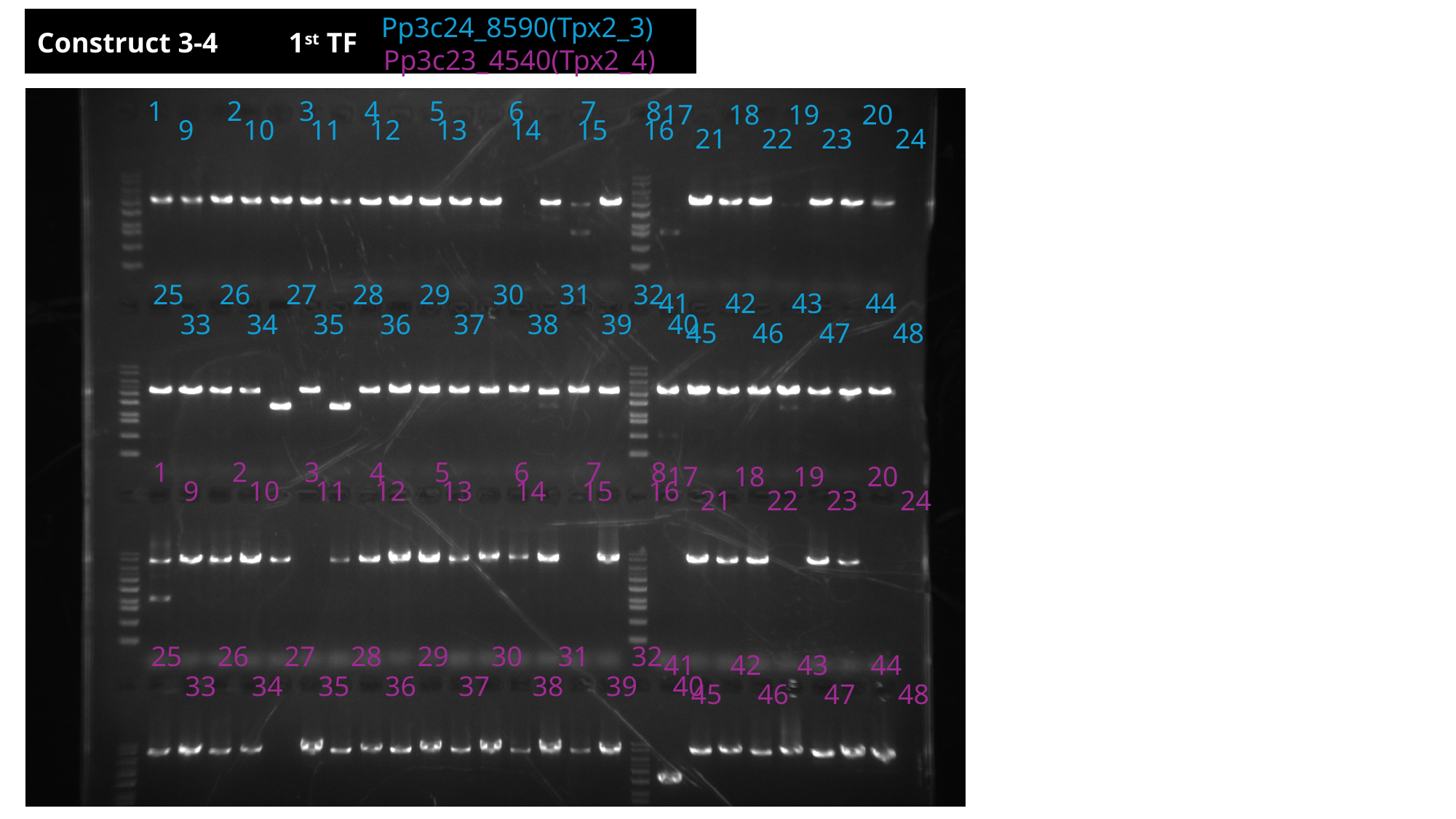

Pp3c24_8590(Tpx2_3)
Construct 3-4 1^st^ TF
Pp3c23_4540(Tpx2_4)
1 2 3 4 5 6 7 8
17 18 19 20
9 10 11 12 13 14 15 16
21 22 23 24
25 26 27 28 29 30 31 32
41 42 43 44
33 34 35 36 37 38 39 40
45 46 47 48
1 2 3 4 5 6 7 8
17 18 19 20
9 10 11 12 13 14 15 16
21 22 23 24
25 26 27 28 29 30 31 32
41 42 43 44
33 34 35 36 37 38 39 40
45 46 47 48

### Slide 15
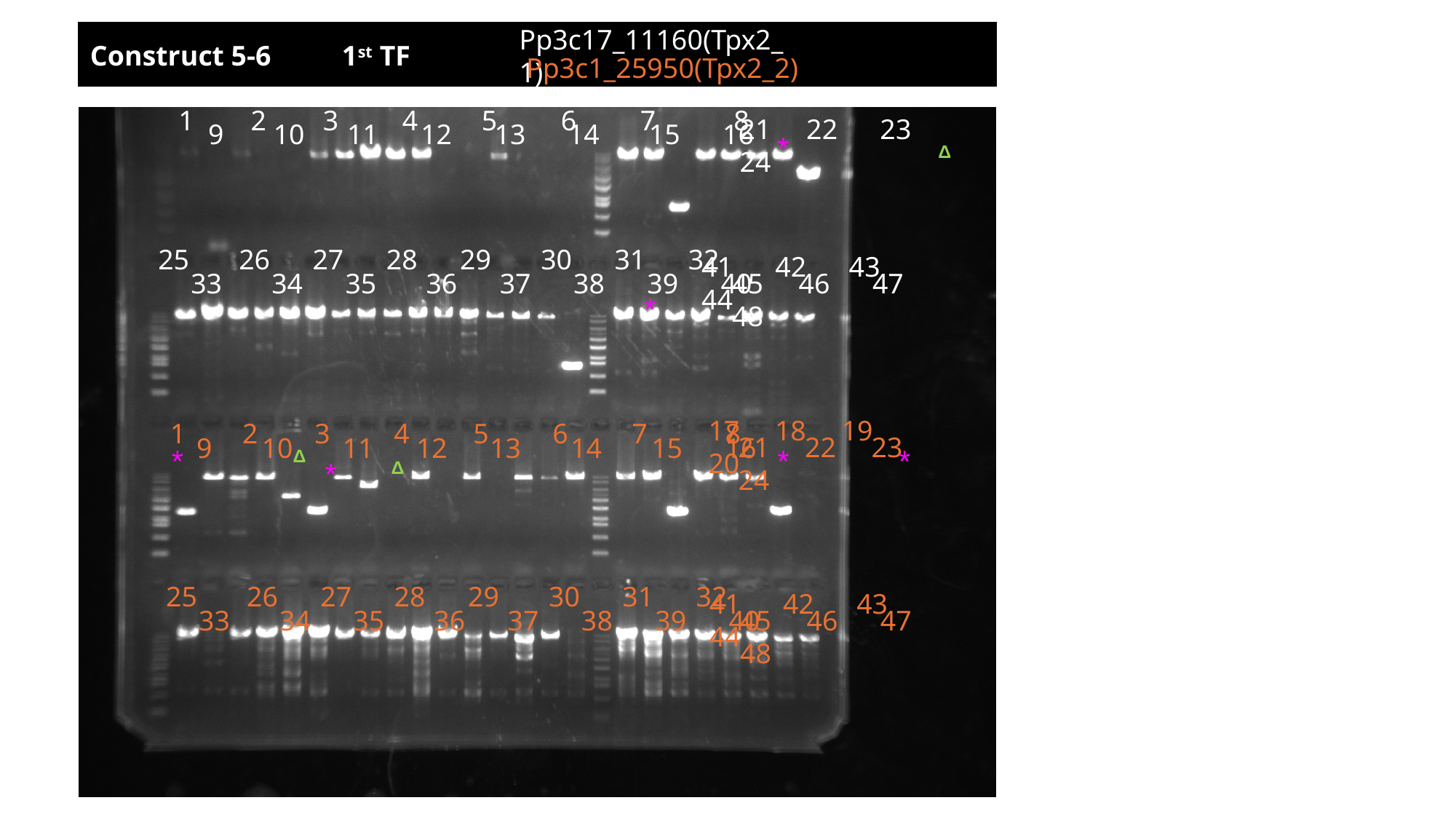

Pp3c17_11160(Tpx2_1)
17 18 19 20
Construct 5-6 1^st^ TF
Pp3c1_25950(Tpx2_2)
1 2 3 4 5 6 7 8
21 22 23 24
9 10 11 12 13 14 15 16
*
Δ
25 26 27 28 29 30 31 32
41 42 43 44
45 46 47 48
33 34 35 36 37 38 39 40
*
17 18 19 20
1 2 3 4 5 6 7 8
21 22 23 24
9 10 11 12 13 14 15 16
*
*
*
Δ
*
Δ
25 26 27 28 29 30 31 32
41 42 43 44
45 46 47 48
33 34 35 36 37 38 39 40

### Slide 16
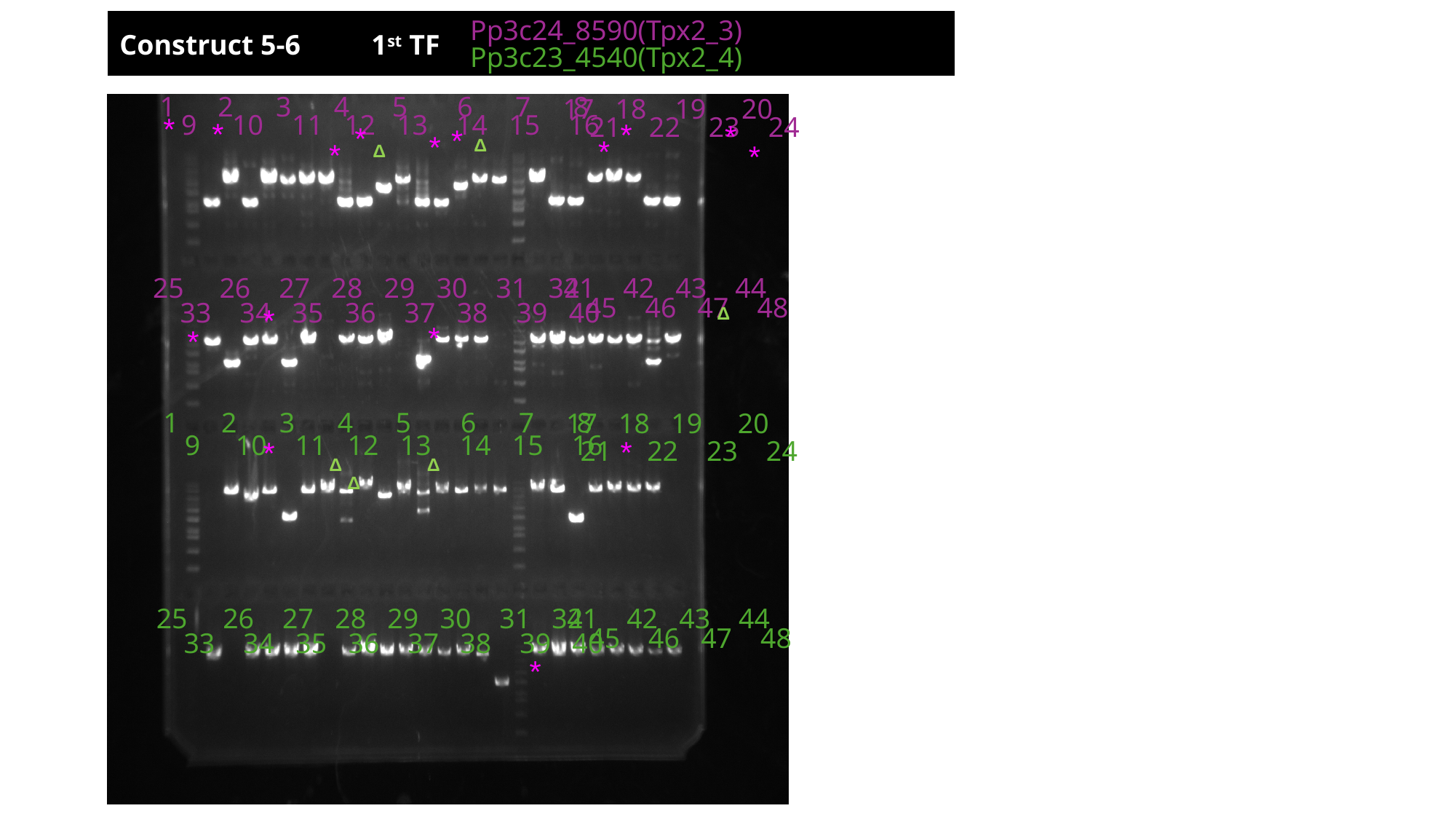

Pp3c24_8590(Tpx2_3)
Construct 5-6 1^st^ TF
Pp3c23_4540(Tpx2_4)
1 2 3 4 5 6 7 8
17 18 19 20
9 10 11 12 13 14 15 16
21 22 23 24
*
*
*
*
*
*
*
*
Δ
*
*
Δ
41 42 43 44
25 26 27 28 29 30 31 32
45 46 47 48
33 34 35 36 37 38 39 40
*
Δ
*
*
1 2 3 4 5 6 7 8
17 18 19 20
9 10 11 12 13 14 15 16
*
21 22 23 24
*
Δ
Δ
Δ
41 42 43 44
25 26 27 28 29 30 31 32
45 46 47 48
33 34 35 36 37 38 39 40
*

### Slide 17
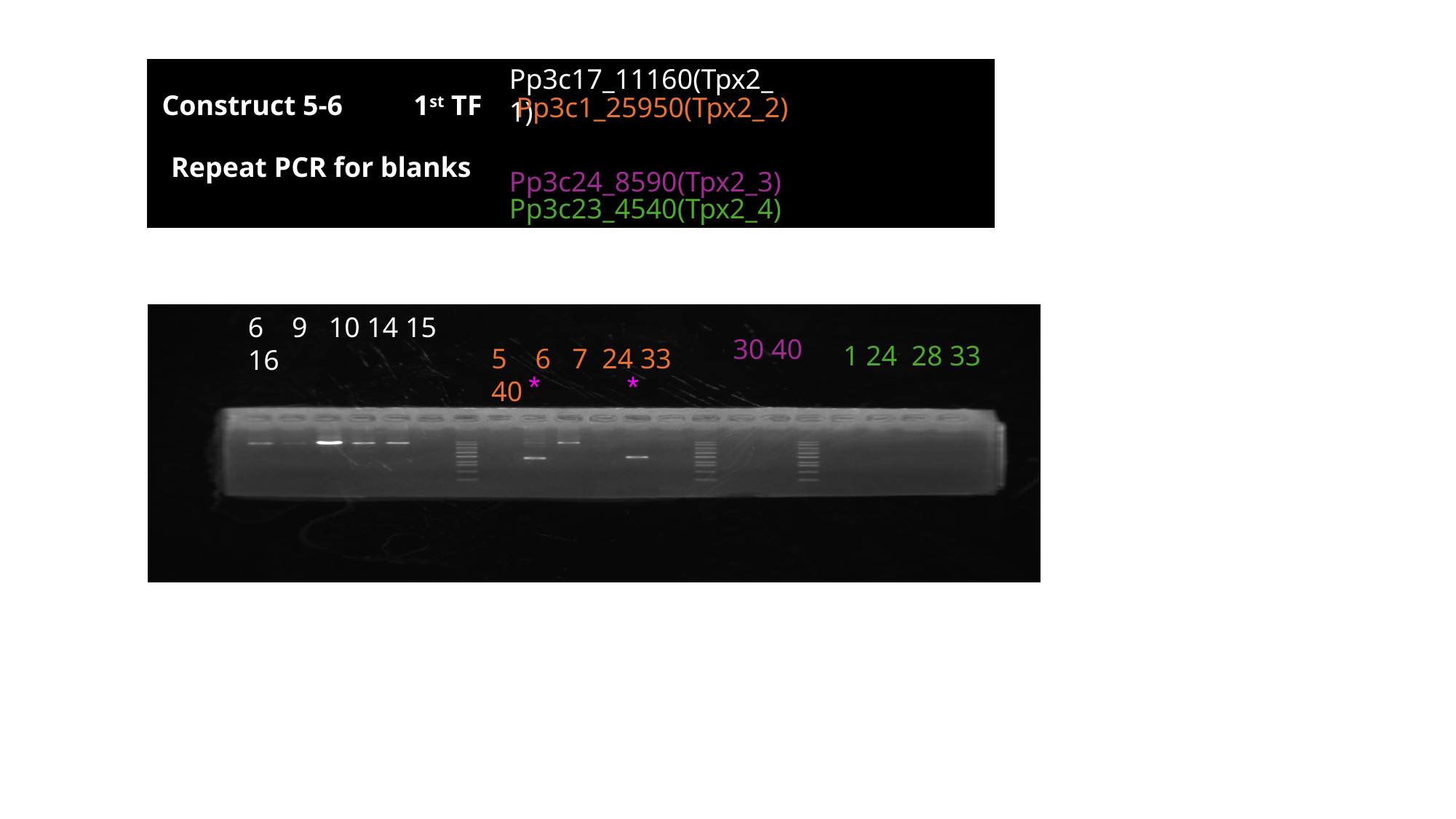

Pp3c17_11160(Tpx2_1)
Construct 5-6 1^st^ TF
Pp3c1_25950(Tpx2_2)
Repeat PCR for blanks
Pp3c24_8590(Tpx2_3)
Pp3c23_4540(Tpx2_4)
6 9 10 14 15 16
30 40
1 24 28 33
5 6 7 24 33 40
*
*

### Slide 18
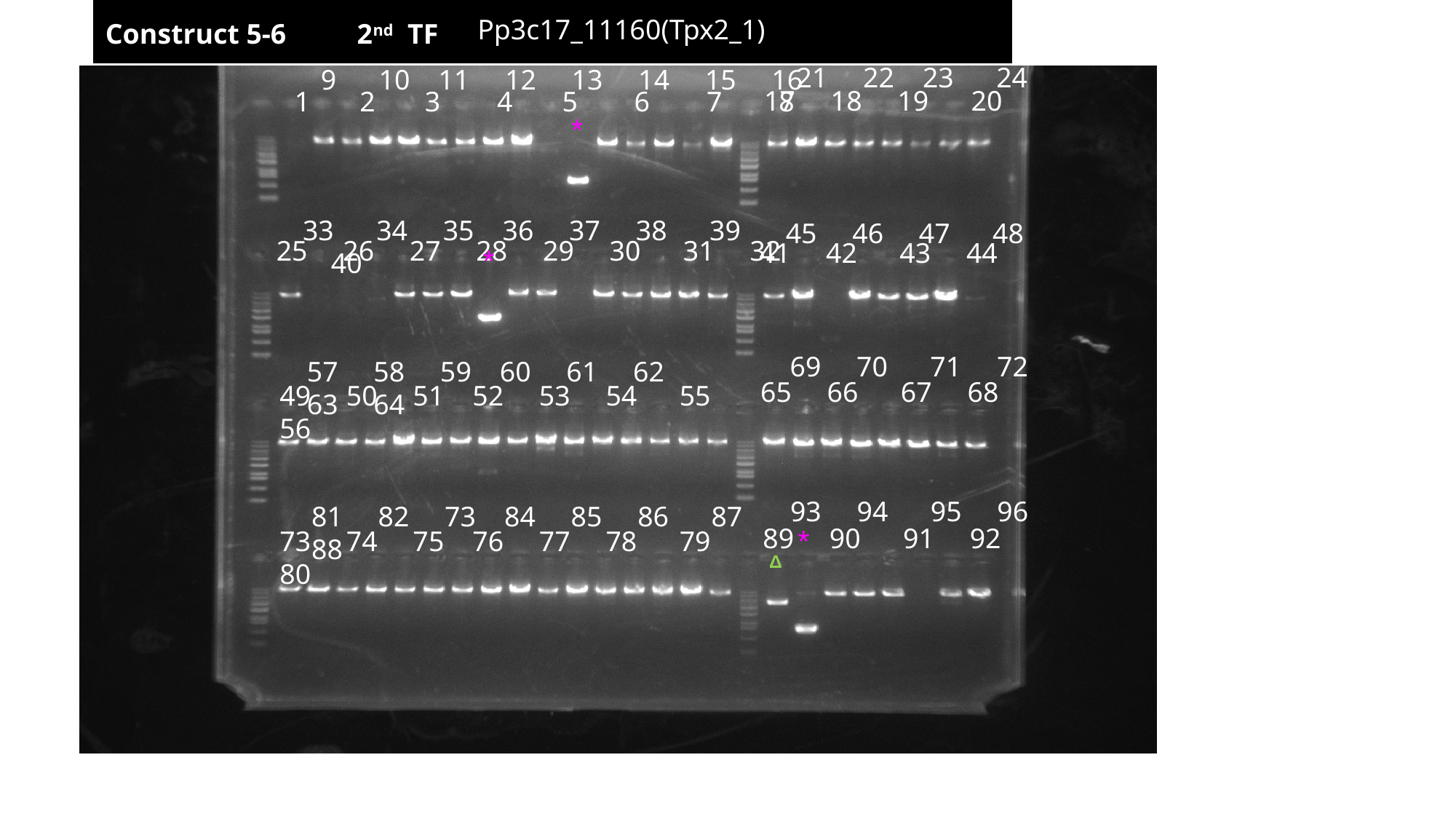

Pp3c17_11160(Tpx2_1)
Construct 5-6 2^nd^ TF
21 22 23 24
9 10 11 12 13 14 15 16
17 18 19 20
1 2 3 4 5 6 7 8
*
33 34 35 36 37 38 39 40
45 46 47 48
25 26 27 28 29 30 31 32
41 42 43 44
*
69 70 71 72
57 58 59 60 61 62 63 64
65 66 67 68
49 50 51 52 53 54 55 56
93 94 95 96
81 82 73 84 85 86 87 88
89 90 91 92
*
73 74 75 76 77 78 79 80
Δ

### Slide 19
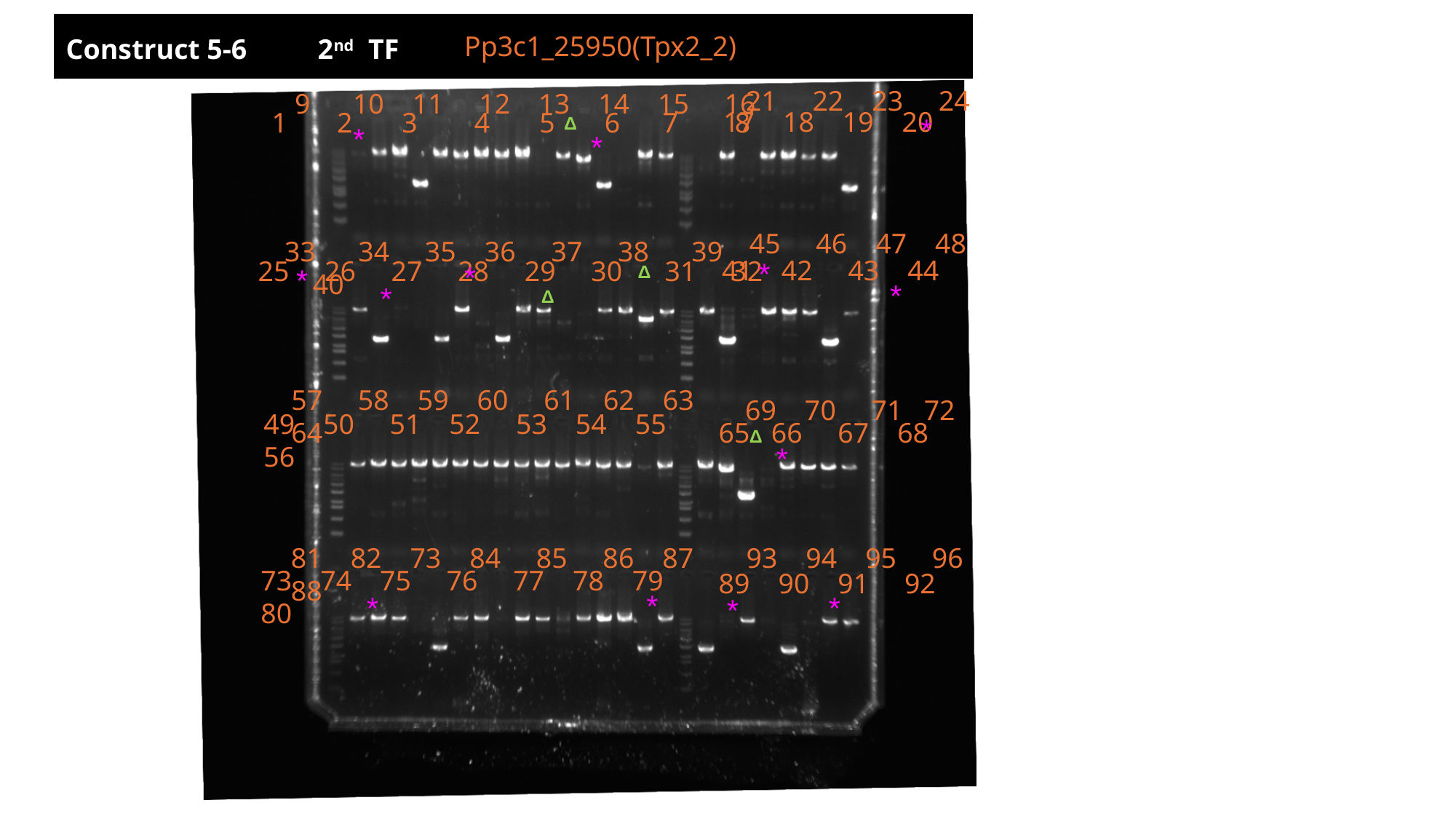

Pp3c1_25950(Tpx2_2)
Construct 5-6 2^nd^ TF
21 22 23 24
9 10 11 12 13 14 15 16
 17 18 19 20
1 2 3 4 5 6 7 8
45 46 47 48
33 34 35 36 37 38 39 40
41 42 43 44
25 26 27 28 29 30 31 32
57 58 59 60 61 62 63 64
69 70 71 72
49 50 51 52 53 54 55 56
65 66 67 68
81 82 73 84 85 86 87 88
93 94 95 96
73 74 75 76 77 78 79 80
89 90 91 92
*
Δ
*
*
*
*
Δ
*
*
*
Δ
Δ
*
*
*
*
*

### Slide 20
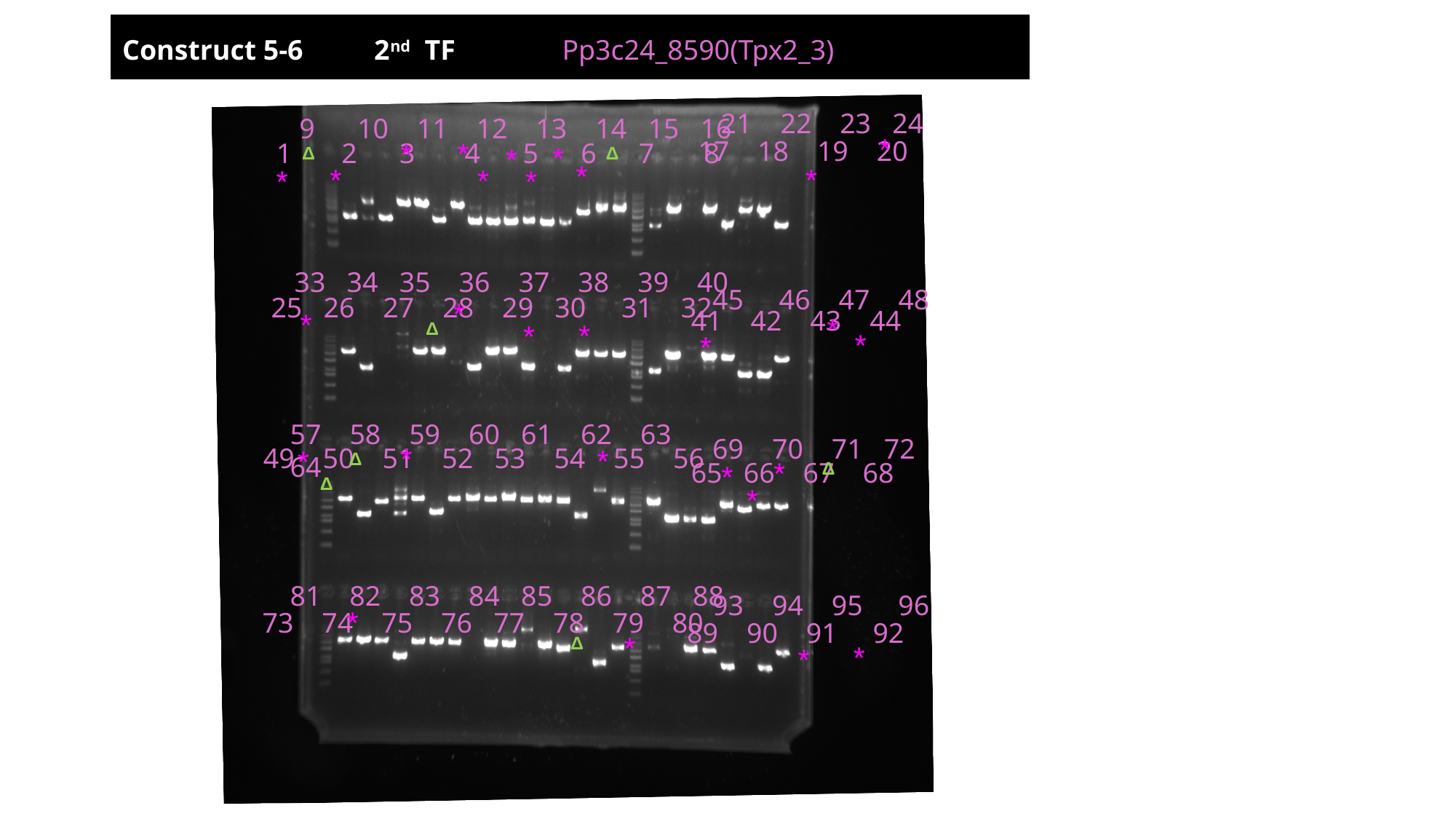

Construct 5-6 2^nd^ TF
Pp3c24_8590(Tpx2_3)
21 22 23 24
9 10 11 12 13 14 15 16
 17 18 19 20
1 2 3 4 5 6 7 8
33 34 35 36 37 38 39 40
45 46 47 48
25 26 27 28 29 30 31 32
41 42 43 44
57 58 59 60 61 62 63 64
69 70 71 72
49 50 51 52 53 54 55 56
65 66 67 68
81 82 83 84 85 86 87 88
93 94 95 96
73 74 75 76 77 78 79 80
89 90 91 92
*
*
*
*
*
Δ
Δ
*
*
*
*
*
*
*
*
*
*
Δ
*
*
*
*
*
*
Δ
*
Δ
*
Δ
*
*
*
Δ
*
*

### Slide 21
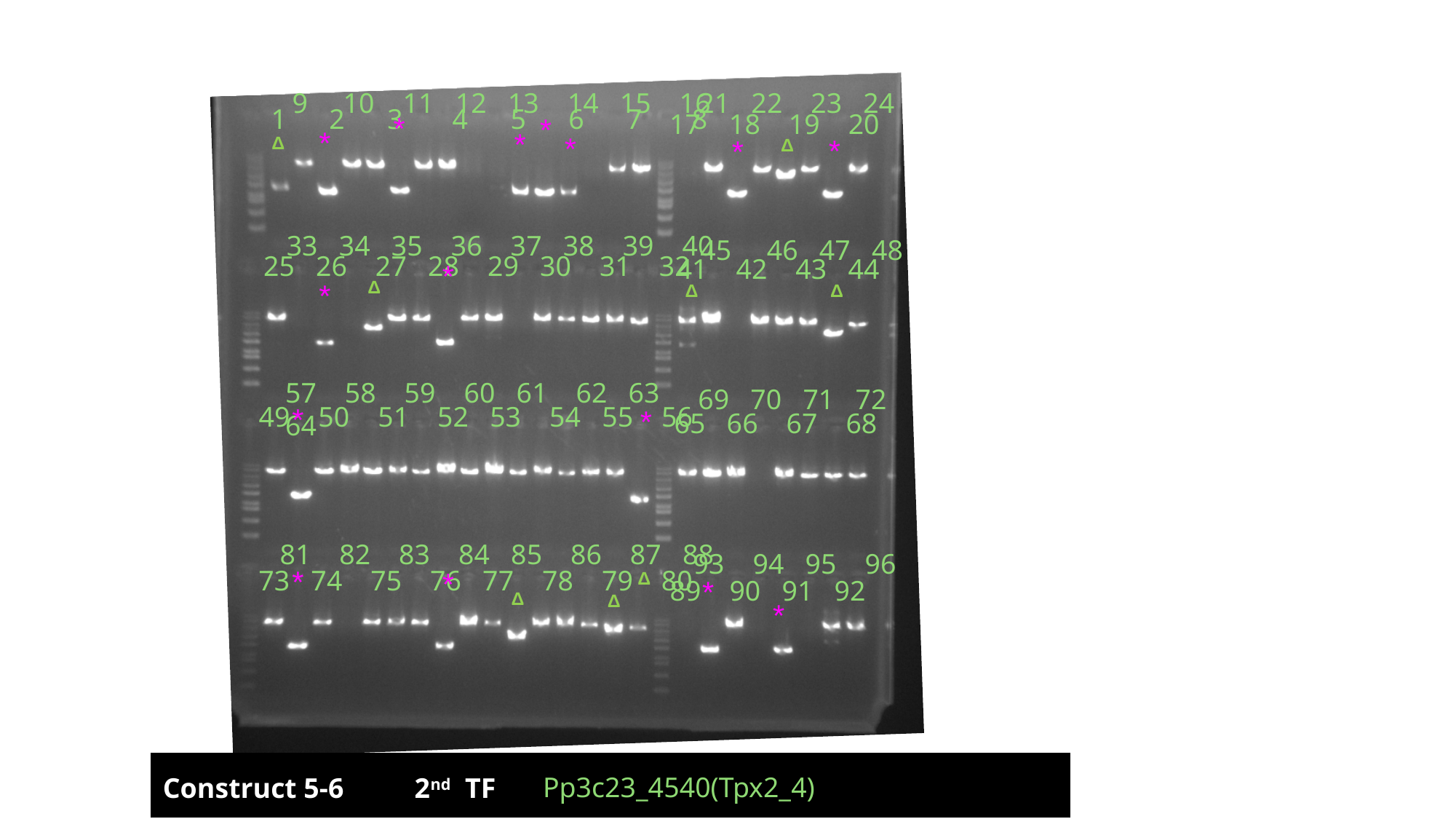

9 10 11 12 13 14 15 16
21 22 23 24
1 2 3 4 5 6 7 8
 17 18 19 20
33 34 35 36 37 38 39 40
45 46 47 48
25 26 27 28 29 30 31 32
41 42 43 44
57 58 59 60 61 62 63 64
69 70 71 72
49 50 51 52 53 54 55 56
65 66 67 68
81 82 83 84 85 86 87 88
93 94 95 96
73 74 75 76 77 78 79 80
89 90 91 92
*
*
*
*
*
*
Δ
*
Δ
*
Δ
*
Δ
Δ
*
*
*
*
Δ
*
Δ
Δ
*
Pp3c23_4540(Tpx2_4)
Construct 5-6 2^nd^ TF

### Slide 22
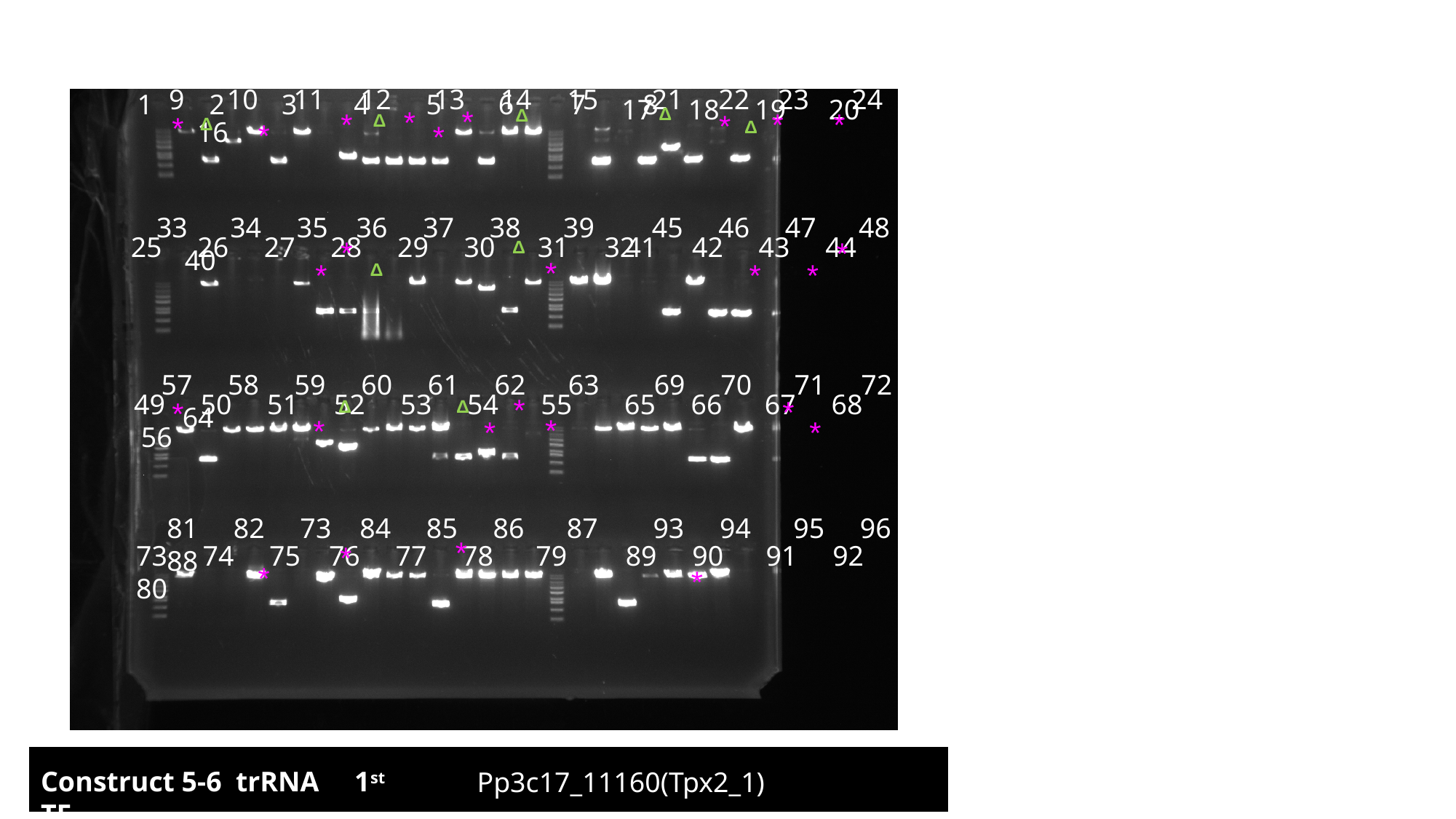

9 10 11 12 13 14 15 16
21 22 23 24
1 2 3 4 5 6 7 8
17 18 19 20
*
Δ
*
Δ
*
*
*
*
*
Δ
Δ
Δ
*
*
33 34 35 36 37 38 39 40
45 46 47 48
25 26 27 28 29 30 31 32
41 42 43 44
*
*
Δ
*
*
*
*
Δ
57 58 59 60 61 62 63 64
69 70 71 72
49 50 51 52 53 54 55 56
65 66 67 68
*
*
*
Δ
Δ
*
*
*
*
81 82 73 84 85 86 87 88
93 94 95 96
*
73 74 75 76 77 78 79 80
*
89 90 91 92
*
*
Construct 5-6 trRNA 1^st^ TF
Pp3c17_11160(Tpx2_1)

### Slide 23
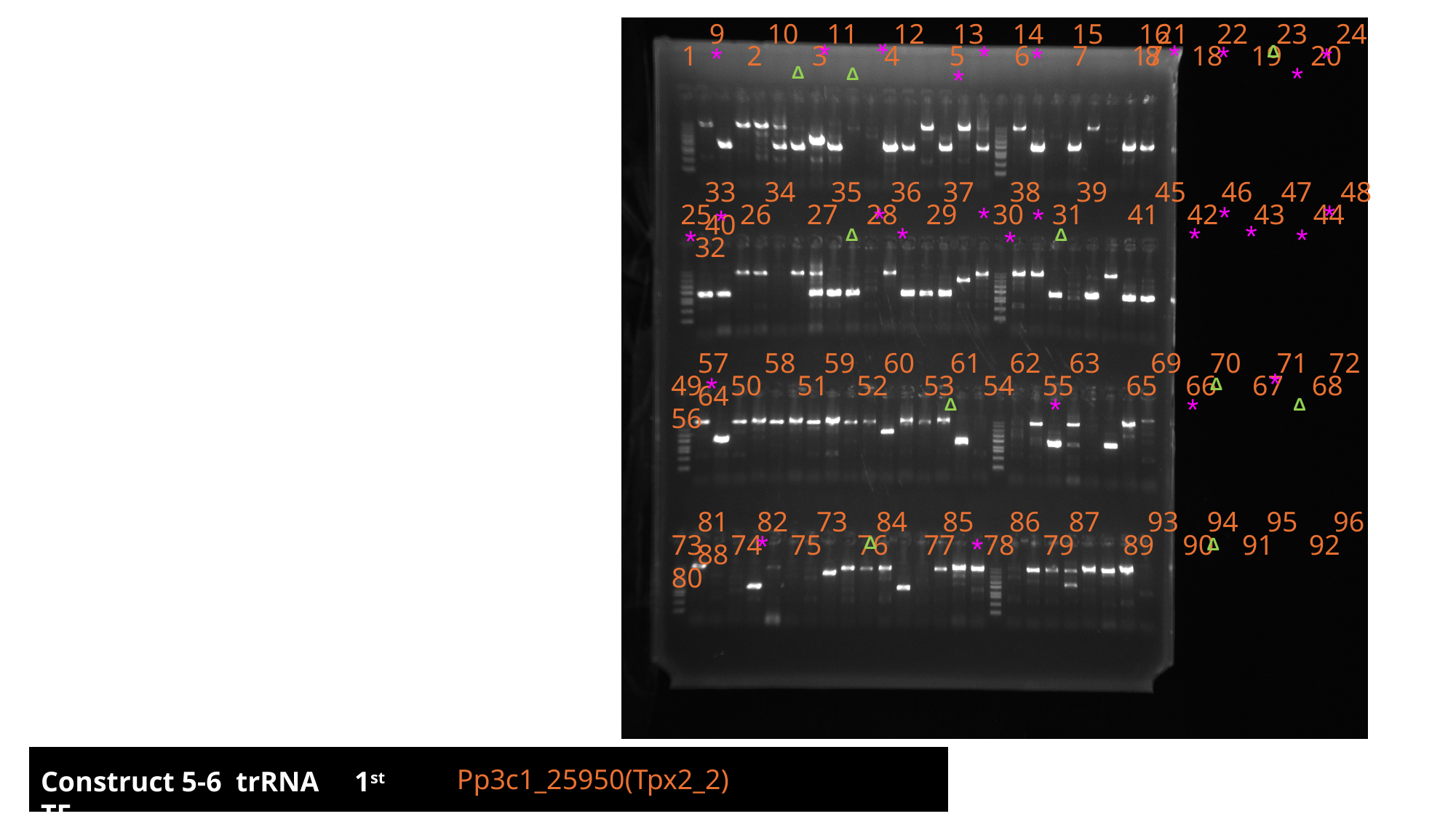

9 10 11 12 13 14 15 16
21 22 23 24
*
*
*
*
*
1 2 3 4 5 6 7 8
 17 18 19 20
*
*
*
Δ
*
*
Δ
Δ
33 34 35 36 37 38 39 40
45 46 47 48
*
25 26 27 28 29 30 31 32
41 42 43 44
*
*
*
*
*
*
*
*
*
*
*
Δ
Δ
57 58 59 60 61 62 63 64
69 70 71 72
*
49 50 51 52 53 54 55 56
65 66 67 68
*
Δ
*
*
Δ
Δ
81 82 73 84 85 86 87 88
93 94 95 96
*
73 74 75 76 77 78 79 80
89 90 91 92
*
Δ
Δ
Pp3c1_25950(Tpx2_2)
Construct 5-6 trRNA 1^st^ TF

### Slide 24
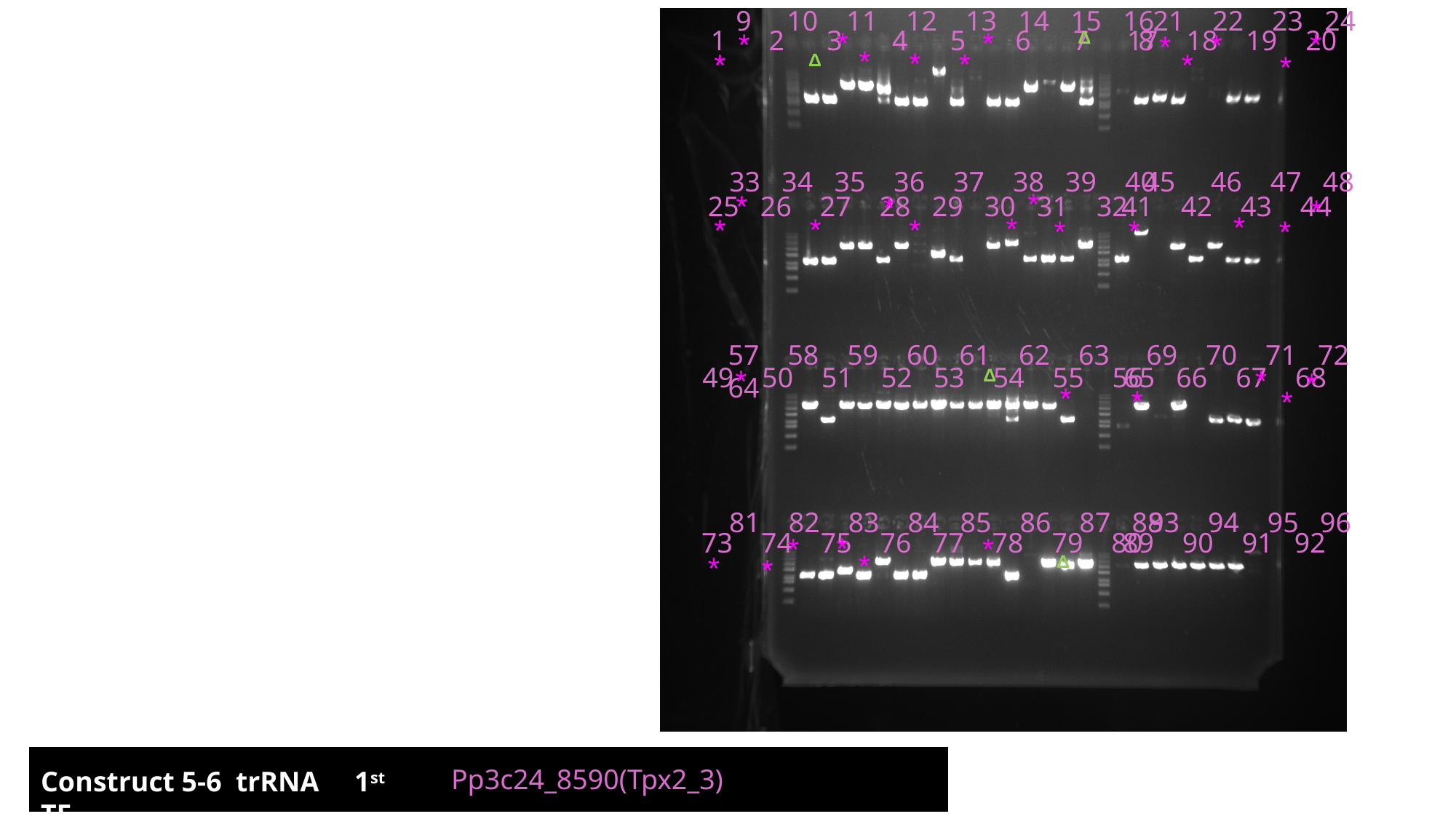

9 10 11 12 13 14 15 16
21 22 23 24
1 2 3 4 5 6 7 8
 17 18 19 20
33 34 35 36 37 38 39 40
45 46 47 48
25 26 27 28 29 30 31 32
41 42 43 44
57 58 59 60 61 62 63 64
69 70 71 72
49 50 51 52 53 54 55 56
65 66 67 68
81 82 83 84 85 86 87 88
93 94 95 96
73 74 75 76 77 78 79 80
89 90 91 92
*
*
*
*
Δ
*
*
*
*
*
*
*
*
Δ
*
*
*
*
*
*
*
*
*
*
*
*
*
*
Δ
*
*
*
*
*
*
*
*
*
Δ
*
Pp3c24_8590(Tpx2_3)
Construct 5-6 trRNA 1^st^ TF

### Slide 25
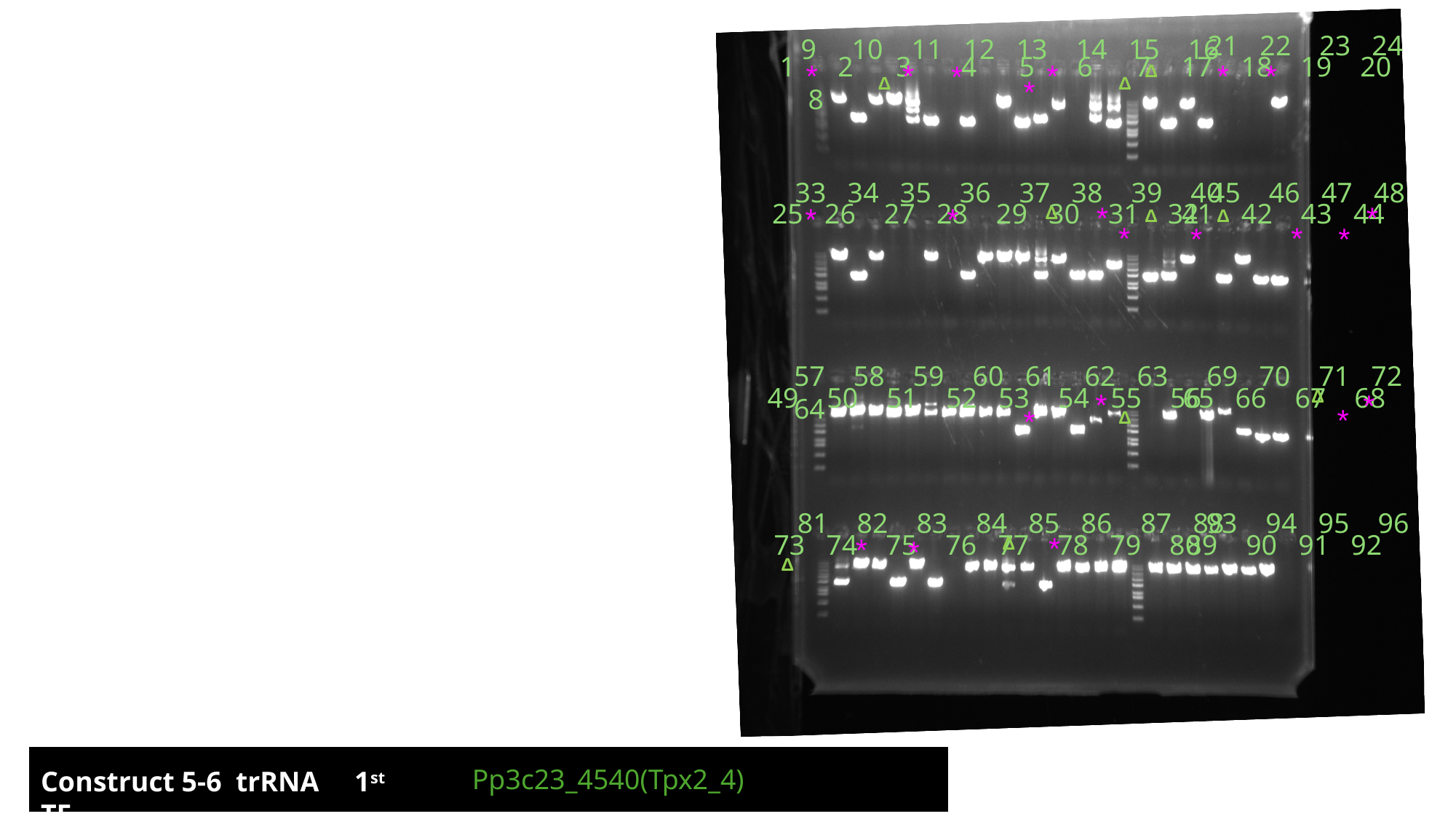

21 22 23 24
9 10 11 12 13 14 15 16
1 2 3 4 5 6 7 8
 17 18 19 20
33 34 35 36 37 38 39 40
45 46 47 48
25 26 27 28 29 30 31 32
41 42 43 44
57 58 59 60 61 62 63 64
69 70 71 72
49 50 51 52 53 54 55 56
65 66 67 68
81 82 83 84 85 86 87 88
93 94 95 96
73 74 75 76 77 78 79 80
89 90 91 92
*
*
*
*
*
*
Δ
Δ
Δ
*
*
*
*
*
Δ
Δ
Δ
*
*
*
*
Δ
*
*
*
*
Δ
*
*
Δ
*
Δ
Pp3c23_4540(Tpx2_4)
Construct 5-6 trRNA 1^st^ TF
